# A direct interaction between CPF and Pol II links RNA 3ʹ-end processing to transcription

**DOI:** 10.1101/2022.07.28.501803

**Authors:** Manuel Carminati, M. Cemre Manav, Dom Bellini, Lori A Passmore

## Abstract

Transcription termination by RNA Polymerase II (Pol II) is linked to RNA 3ʹ-end processing by the cleavage and polyadenylation factor (CPF). CPF contains endonuclease, poly(A) polymerase and protein phosphatase activities that cleave and polyadenylate the pre-mRNA, and dephosphorylate Pol II to control transcription. Exactly how the RNA 3ʹ-end processing machinery is coupled to transcription remains unclear. Here, we combine *in vitro* reconstitution, electron cryo-microscopy and X-ray crystallography to show that CPF physically and functionally interacts with Pol II. Surprisingly, CPF-mediated dephosphorylation promotes the formation of a Pol II stalk-to-stalk homodimer. This dimer is compatible with transcription (unlike other Pol II dimers) but not with the binding of transcription elongation factors. We show that the Ref2 subunit of CPF contributes directly to the interaction with Pol II and also regulates the Glc7/PP1 CPF phosphatase. We hypothesize that Pol II dimerization may play a role in gene looping and may provide a mechanistic basis for the allosteric model of transcription termination.

## Introduction

Eukaryotic mRNAs must be processed before they can be exported from the nucleus and translated into proteins. This includes capping at their 5ʹ-end, splicing, and 3ʹ-cleavage and polyadenylation. Crosstalk between these processes promotes coordinated and timely recruitment of transcription factors, 5ʹ-capping proteins, the spliceosome and the 3ʹ-end processing machinery to Pol II and nascent pre-mRNAs (Hirose and Manley, 1998; Hirose et al., 1999; Mischo and Proudfoot, 2013). Together, this enhances the efficiency and accuracy of pre-messenger RNA (pre-mRNA) maturation (Bentley, 2014; Herzel et al., 2017).

Transcription and RNA processing are coordinated by regulated phosphorylation of the C-terminal domain (CTD) of Rpb1, the largest subunit of Pol II. The CTD is comprised of 26 heptapeptide repeats in yeast (52 in humans) with a consensus sequence of Y_1_S_2_P_3_T_4_S_5_P_6_S_7_ that is differentially phosphorylated throughout the transcription cycle. The CTD is unphosphorylated when Pol II is loaded onto promoters during transcription initiation (Zhao et al., 1999). Phosphorylation of CTD Ser5 (P-Ser5) peaks early in transcription, and plays a role in recruiting transcription elongation factors, 5ʹ-capping proteins, splicing factors (Ghosh et al., 2011; Komarnitsky et al., 2000; Nojima et al., 2018) and, on short non-coding RNAs, termination factors (Vasiljeva et al., 2008). When Pol II engages in productive elongation, P-Ser5 levels decrease and phosphorylation of Tyr1 and Ser2 residues (P-Tyr1 and P-Ser2) accumulates. P-Tyr1 prevents premature termination of transcription (Mayer et al., 2012; Porrua and Libri, 2015). P-Tyr1 is dephosphorylated after transcription of the polyadenylation signal (PAS), unmasking the ability of P-Ser2 to recruit transcription termination factors (Licatalosi et al., 2002; Mayer *et al*., 2012).

mRNA 3ʹ-end processing is carried out by the Cleavage and Polyadenylation Factor (CPF in yeast; CPSF in humans). CPF is an ∼1 MDa complex containing 14 subunits that are organized into three enzymatic modules (Casañal et al., 2017). The 3-subunit nuclease module contains the Ysh1 (CPSF73) endonuclease that cleaves nascent RNA at specific sites (Chanfreau et al., 1996). The 5-subunit polymerase module contains Pap1 (PAP), which adds a poly(A) tail onto the new free 3ʹ-end (Kumar et al., 2019; Sun et al., 2020). The 6-subunit phosphatase module harbors Ssu72 (SSU72) and Glc7 (the yeast PP1 phosphatase). Ssu72 is a P-Ser5 phosphatase that plays roles in transcription elongation (Singh and Hampsey, 2007; Xiang et al., 2010) and gene looping, where the promoter and terminator of a gene are juxtaposed to facilitate Pol II recycling and regulate transcription directionality (Allepuz-Fuster et al., 2019; Tan-Wong et al., 2012). Glc7 dephosphorylates P-Tyr1 of the CTD to elicit transcription termination (Schreieck et al., 2014). The human and fission yeast orthologues of Glc7, PP1 and Dis2 respectively, also promote transcription termination through dephosphorylation of the elongation factor Spt5 (Cortazar et al., 2019; Kecman et al., 2018). While CPF processes the 3ʹ-ends of protein-coding transcripts, the related APT (<u>A</u>ssociated with <u>Pt</u>a1) complex is required for transcription of non-coding snRNAs and snoRNAs by Pol II in yeast (Lidschreiber et al., 2018; Nedea et al., 2008). APT contains all six subunits of the phosphatase module as well as an additional subunit, Syc1.

Transcription and mRNA 3ʹ-end processing are intimately coupled across eukaryotes (Connelly and Manley, 1988; Logan et al., 1987). CPF interacts with the transcription initiation complex at promoters, remains associated with Pol II throughout transcription, and is required for transcription termination (Baejen et al., 2017; Dantonel et al., 1997; Zhao *et al*., 1999). There are two prevalent, non-exclusive models for transcription termination. First, in the torpedo model, pre-mRNA cleavage by CPF is required to allow access of the 5ʹ–3ʹ ‘torpedo’ exonuclease that releases Pol II from DNA (Kim et al., 2004; West et al., 2004). In the allosteric model, transcription of the PAS triggers an undefined change in Pol II that renders it competent for termination (Logan *et al*., 1987; Richard and Manley, 2009). CPF-mediated dephosphorylation of Pol II likely contributes to the allosteric model (Schreieck *et al*., 2014).

Despite the coupling between 3ʹ-end processing and transcription, it remains unclear how or whether CPF binds transcribing Pol II. Moreover, it is unknown how Pol II affects 3ʹ-end processing. Here, we report that CPF and APT directly bind Pol II. This interaction is partly mediated via the Ref2 subunit, which also regulates the CPF phosphatase Glc7. We show that dephosphorylation of the CTD promotes the formation of Pol II homodimers that are structurally incompatible with transcription elongation. Together, this establishes that transcription and 3ʹ-end processing are physically connected and suggests that the phosphatase activity of CPF may regulate transcription termination by controlling the oligomerization state of Pol II.

## Results

### CPF and APT interact directly with Pol II

Since transcription and RNA 3ʹ-end processing are functionally coupled by Pol II CTD dephosphorylation (Schreieck *et al*., 2014; Xiang *et al*., 2010), we tested whether Pol II interacts directly with CPF or APT in a fully defined *in vitro* system. We purified an affinity-tagged Pol II from yeast, and recombinant yeast CPF, CPF-core (polymerase and nuclease modules), phosphatase module, and APT from insect cells after baculovirus-based overexpression (Figure S1A, lanes 1-4). We then performed CPF and APT pull-down assays using immobilized Pol II. Interestingly, both CPF and APT bound to Pol II *in vitro* (Figure 1A). Moreover, we found that the phosphatase module of CPF is necessary and sufficient for the interaction (Figure S1B).

**Figure 1.**
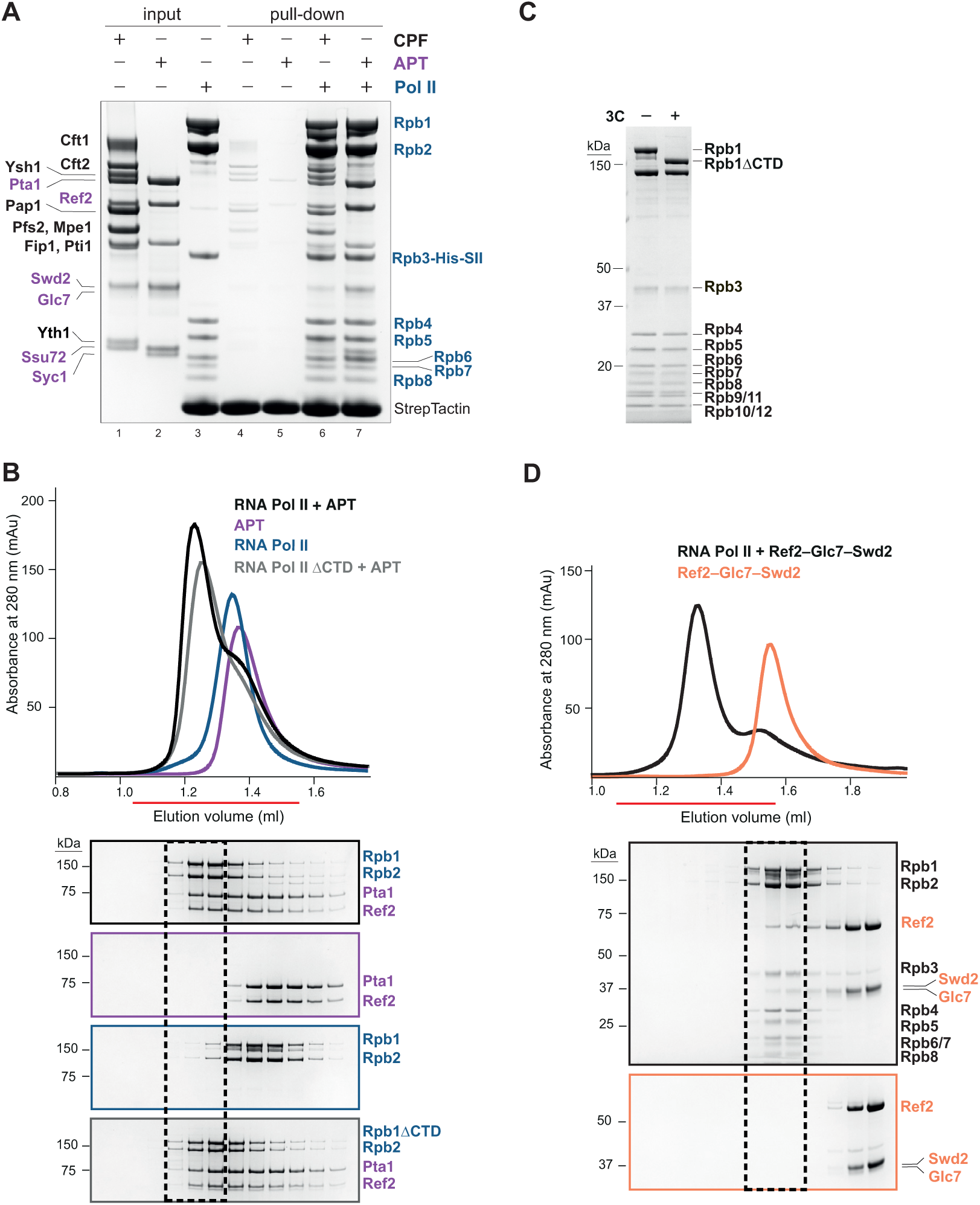
CPF and APT interact directly with Pol II. (**A**) Pulldown assay of CPF or APT using StrepII (SII)-tagged Pol II (Rpb3-His-SII) immobilized on StrepTactin beads. Input and bound proteins were analyzed on SDS-PAGE. APT subunits are labelled in purple and the remaining CPF subunits are labelled in black (left); Pol II subunits are labelled in blue (right). The experiment was repeated twice. (**B**) (Top) Analytical size exclusion chromatography profiles of Pol II (blue) and APT (purple), loaded separately or after incubation (black). APT in complex with Pol II- CTD is in grey. (Bottom) SDS-PAGE (cropped) of the fractions indicated by a red line. The black dashed box indicates the migration position of the Pol II–APT complex. Gels are outlined according to colors of chromatograms. Uncropped gels are in Supplementary Data 1A. (**C**) SDS-PAGE showing 3C protease cleavage of the Rpb1 CTD to make Poll II- CTD. (**D**) Analytical size exclusion chromatography experiment as in **B**, of Pol II incubated with the Ref2–Glc7–Swd2 subcomplex of APT. The fractions indicated by a red line were analyzed by the SDS-PAGE below.

As the six subunits of the phosphatase module are also found within APT, we reasoned that both complexes are likely to interact with Pol II in a similar manner. We therefore focused our subsequent analyses on APT, which is more stable and less prone to degradation than the phosphatase module *in vitro*, likely because it is a complete complex rather than a subcomplex of CPF. A complex of APT and Pol II co-eluted on analytical size exclusion chromatography showing that they form a stable complex in solution (Figure 1B). In summary, these data show that APT and the phosphatase module of CPF directly bind Pol II.

### APT interacts with the core of Pol II mainly via Ref2–Glc7–Swd2

Next, we investigated which region of Pol II interacts with APT. Since the CTD of Pol II is the substrate of APT, it could be involved in their interaction. The CTD is essential for viability in yeast (West and Corden, 1995) and therefore, we used CRISPR-Cas9 to introduce a 3C protease cleavage site into Rpb1 between the core of the protein and the CTD. This allowed removal of the CTD after Pol II purification (Figure 1C). We incubated Pol II-ΔCTD with APT and found that they still co-elute on size exclusion chromatography (Figure 1B), demonstrating that they interact even in the absence of the CTD.

Previous data showed that CPF promotes the formation of gene loops in yeast through an interaction between Ssu72 and Rpb4, a subunit of the Pol II stalk (Allepuz-Fuster *et al*., 2019). To investigate whether APT interacts with Pol II via the stalk (comprised of Rpb4 and Rpb7), we purified a stalk-less version of Pol II (Pol II-Δstalk) from an *RPB4* deletion strain (Allepuz-Fuster et al., 2014; Allepuz-Fuster *et al*., 2019). Stalk-less Pol II also retained the ability to interact with APT on size exclusion chromatography (Figure S1C). Thus, the primary binding site for APT is within the core of Pol II, not in the CTD or stalk.

Next, to define which region of APT interacts with Pol II, we purified two subcomplexes of APT that we had previously identified by native mass spectrometry (Ref2–Glc7–Swd2 and Pta1–Pti1–Ssu72–Syc1 respectively; Figure S1A, lanes 5-6) (Casañal *et al*., 2017). In a pull-down assay, both the Glc7- and Ssu72-containing subcomplexes interact with Pol II (Figure S1D). The Glc7-subcomplex pulled-down more Pol II so we focused on it. In size exclusion chromatography, the migration position of the Glc7-subcomplex shifts after incubation with Pol II, confirming a direct interaction between this subcomplex and the polymerase (Figure 1D).

Together, our data show that neither the CTD nor the stalk are required for APT to bind Pol II. Moreover, we found that the Ref2–Glc7–Swd2 subcomplex makes a major contribution towards the interaction of APT with Pol II.

### APT binds Pol II next to the RNA exit channel

To gain insight into the functional interplay between Pol II and the 3ʹ-end processing machinery, we tested whether *in vitro* transcription by Pol II is influenced by the presence of APT or CPF in a promoter-independent RNA elongation assay. We made an artificial transcription bubble using a model pre-mRNA substrate (part of the yeast *CYC1* transcript), which is elongated by Pol II in the presence of ribonucleotides. This showed that basal transcription by Pol II is compatible with the presence of APT and CPF (Figure S2A).

The Pol II CTD has been proposed to stimulate 3ʹ-cleavage (Hirose and Manley, 1998). Thus, we tested this in a fully reconstituted system. We assembled Pol II on a transcription bubble containing a longer version (259 nt) of the model pre-mRNA substrate, mimicking transcription of an RNA with an upstream cleavage site. We carried out cleavage assays in the presence of CPF, as well as the accessory factors CF IA and CF IB. Our results show that CPF mediates efficient and specific cleavage of pre-mRNA in the presence of Pol II but its activity is neither stimulated nor inhibited (Figure S2B).

To understand whether there are other functional consequences of physically connecting the RNA 3ʹ-end processing machinery to Pol II, we examined the structure of APT in complex with Pol II by electron cryo-microscopy (cryoEM). Thus, we assembled a complex between purified APT and Pol II loaded onto a DNA–RNA scaffold, which mimics a transcription elongation bubble. Because APT specifically plays a role in transcription of non-coding transcripts, we assembled a transcription bubble using a small nucleolar RNA (snR47) which had been previously linked to APT activity (Lidschreiber *et al*., 2018) (Figure S2C). The RNA 3ʹ-end processing machinery is generally unstable in cryoEM (Hill et al., 2019), so we prepared cryo-EM grids after gradient centrifugation coupled to glutaraldehyde crosslinking (GraFix) (Kastner et al., 2008) to stabilize the complex (Figure S3A).

3D-reconstructions of the transcribing complex show Pol II with additional density next to the RNA exit channel and the stalk (Figure 2A; Figure S3B-C). The additional density is poorly resolved, indicating that APT (and presumably CPF) may be flexibly positioned on Pol II. This binding site of APT is in agreement with a previously-reported interaction of Ssu72 with the stalk protein Rpb4 (Allepuz-Fuster *et al*., 2019), which also occupies a position proximal to the nascent RNA exit channel. Together, these data show that the RNA 3ʹ-end processing machinery is positioned on Pol II so that it can monitor the nascent transcript as it emerges from the RNA exit channel. Flexibility would allow it to access RNAs with diverse lengths and structures.

**Figure 2.**
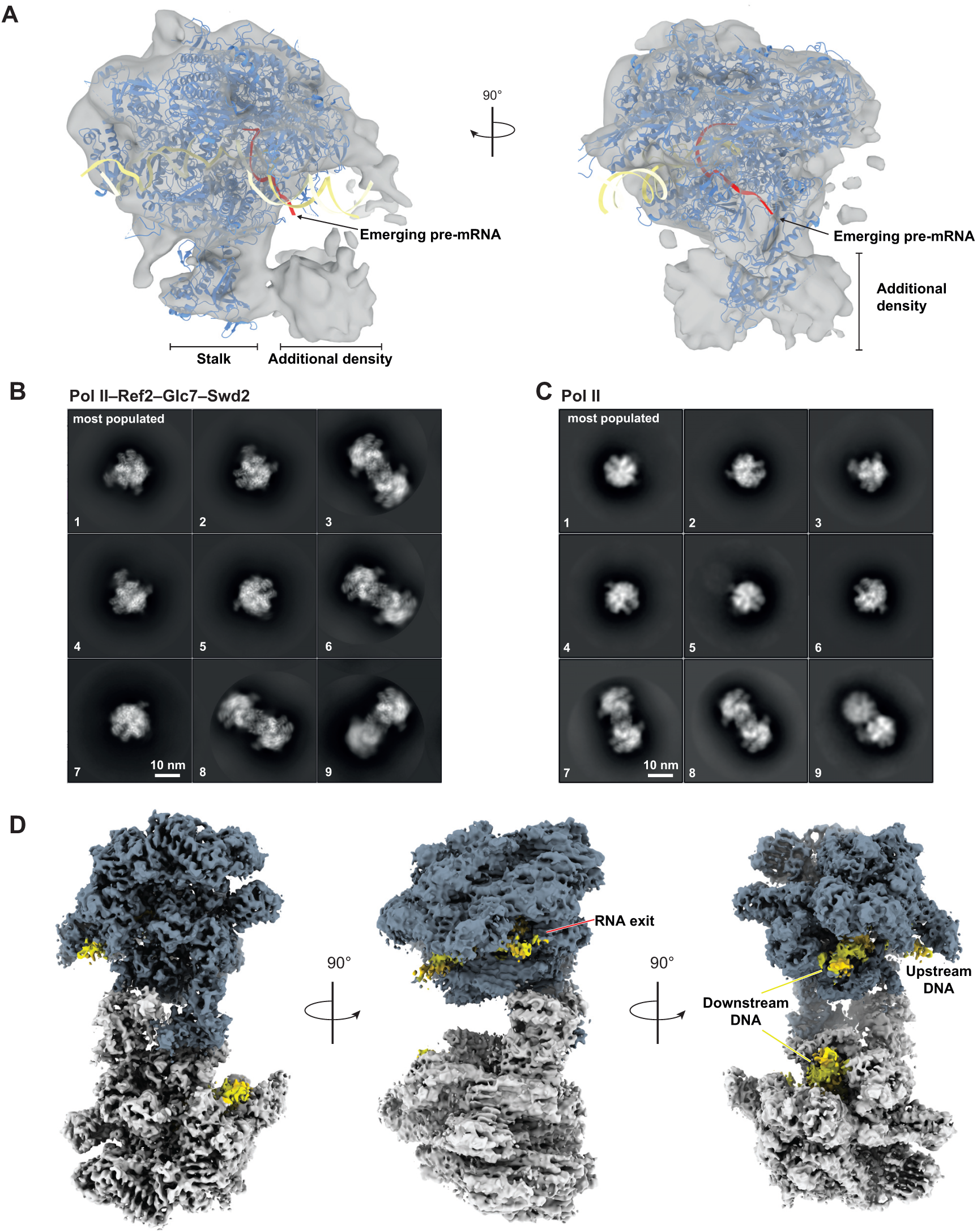
CryoEM analysis of Pol II reveals Pol II homodimers in the presence of Ref2–Glc7–Swd2. (**A**) 3D reconstruction of Pol II–APT (grey transparent surface) with a model of Pol II (PDB 5C4X; blue ribbon) rigid-body fit into the map. DNA is in yellow, and RNA is in red. Density not accounted for by Pol II is indicated. Arrows point at the RNA exit channel. (**B**) Selected 2D class-averages of DNA–RNA-loaded Pol II with Ref2–Glc7–Swd2 from 850,000 particles showing Pol II monomers and dimers (see also Figure S4B-C). Classes are ordered by the number of particles in each class, from the most populated (1) to the least populated (9). (**C**) Selected 2D class-averages of DNA–RNA-loaded Pol II from 400,000 particles, ordered as in panel **B**. (**D**) Composite map of the Pol II dimer after signal subtraction and focused 3D-refinement of the individual monomers (overall resolution 3.6 Å). One monomer is colored in blue; the other is in grey. The DNA-RNA scaffold is in yellow. See Figure S3 and 5 for the EM processing pipelines, and Figure S4E for the map of intact dimeric Pol II before signal subtraction.

### Ref2–Glc7–Swd2 promotes the formation of Pol II homodimers

To further investigate the association of the 3ʹ-end processing machinery with Pol II, we next assembled the smaller Ref2–Glc7–Swd2 subcomplex of APT with Pol II loaded onto a DNA–RNA hybrid (Figure S4A). We vitrified the sample in native conditions (without crosslinkers) but in the presence of detergents to minimize the orientation bias known to affect Pol II (Davis et al., 2002; Ehara *et al*., 2017; Lahiri et al., 2019). In 2D-class averages, we did not identify any obvious density for Ref2–Glc7–Swd2 (Figure 2B and Figure S4B-C). Unexpectedly, we observed many Pol II homodimers in this dataset. Pol II dimers were also present in the absence of Ref2–Glc7–Swd2 but at a much lower proportion (Figure 2C). Dimers were not evident in our crosslinked samples, but they have been observed in previous studies (Aibara et al., 2021). Crosslinking may therefore antagonize dimer formation.

We obtained a 3D map of the Pol II homodimer at 4.6 Å resolution (Figure S4D-E and Table 1). Compared to the structure of Pol II alone, there was no additional density that could account for Ref2–Glc7–Swd2. To further improve the Pol II homodimer map, we carried out focused 3D refinement on each monomer after signal subtraction by masking one monomer at a time (Figure S5 and 6). We then combined the two resulting maps, yielding a composite map of the intact Pol II dimer at an overall resolution of 3.6 Å (Figure 2D and Table 1).

**Table 1.** Cryo-EM data collection and processing.

|  | Pol II dimer<br>(EMDB-15358) | Pol II dimer<br>focused<br>refinement on<br>monomer 1<br>(EMDB-15359) | Pol II dimer<br>focused<br>refinement on<br>monomer 2<br>(EMDB-15360) |
| --- | --- | --- | --- |
| **Data collection and processing** |  |  |  |
| Magnification * UCam | 105,000 x | 105,000 x | 105,000 x |
| LMB Krios3 | 105,000 x | 105,000 x | 105,000 x |
| LMB Krios1 | 75,000 x | 75,000 x | 75,000 x |
| Voltage (kV) | 300 | 300 | 300 |
| Electron exposure (e-/Å<sup>2</sup>) | 40 | 40 | 40 |
| Defocus range (µm) | 1.5-2.7 | 1.5-2.7 | 1.5-2.7 |
| Pixel size (Å)* UCam | 0.83 | 0.83 | 0.83 |
| LMB Krios3 | 0.86 | 0.86 | 0.86 |
| LMB Krios1 | 1.04 | 1.04 | 1.04 |
| Symmetry imposed | C1 | C1 | C1 |
| Initial particle images (no.) | 5,800,000 | 5,800,000 | 5,800,000 |
| Final particle images (no.) | 151,000 | 151,000 | 151,000 |
| Map resolution (Å) | 4.6 | 3.6 | 3.6 |
| FSC threshold | 0.143 | 0.143 | 0.143 |
| Map resolution range (Å) | 3.8-17.5 | 3.3-11.8 | 3.3-9.0 |
| Map sharpening *B* factor (Å<sup>2</sup>) | -100 | -50 | -50 |

\* Datasets collected on three different detectors. Magnification and pixel size refer respectively to K3-Gatan, University of Cambridge; K3-Gatan, Krios III, MRC-LMB; Falcon3, Krios I, MRC-LMB.

### Pol II dimerizes via the stalk

We fitted two copies of a crystal structure of monomeric Pol II (Barnes *et al*., 2015) into the composite map of the Pol II homodimer (Figure 3A). Overall, these models agree well with the map within the Pol II core, including the DNA–RNA hybrid. There were some deviations between the map and the models for the stalk regions (Figure S7A), suggesting that the stalk is somewhat rearranged upon dimer formation.

**Figure 3.**
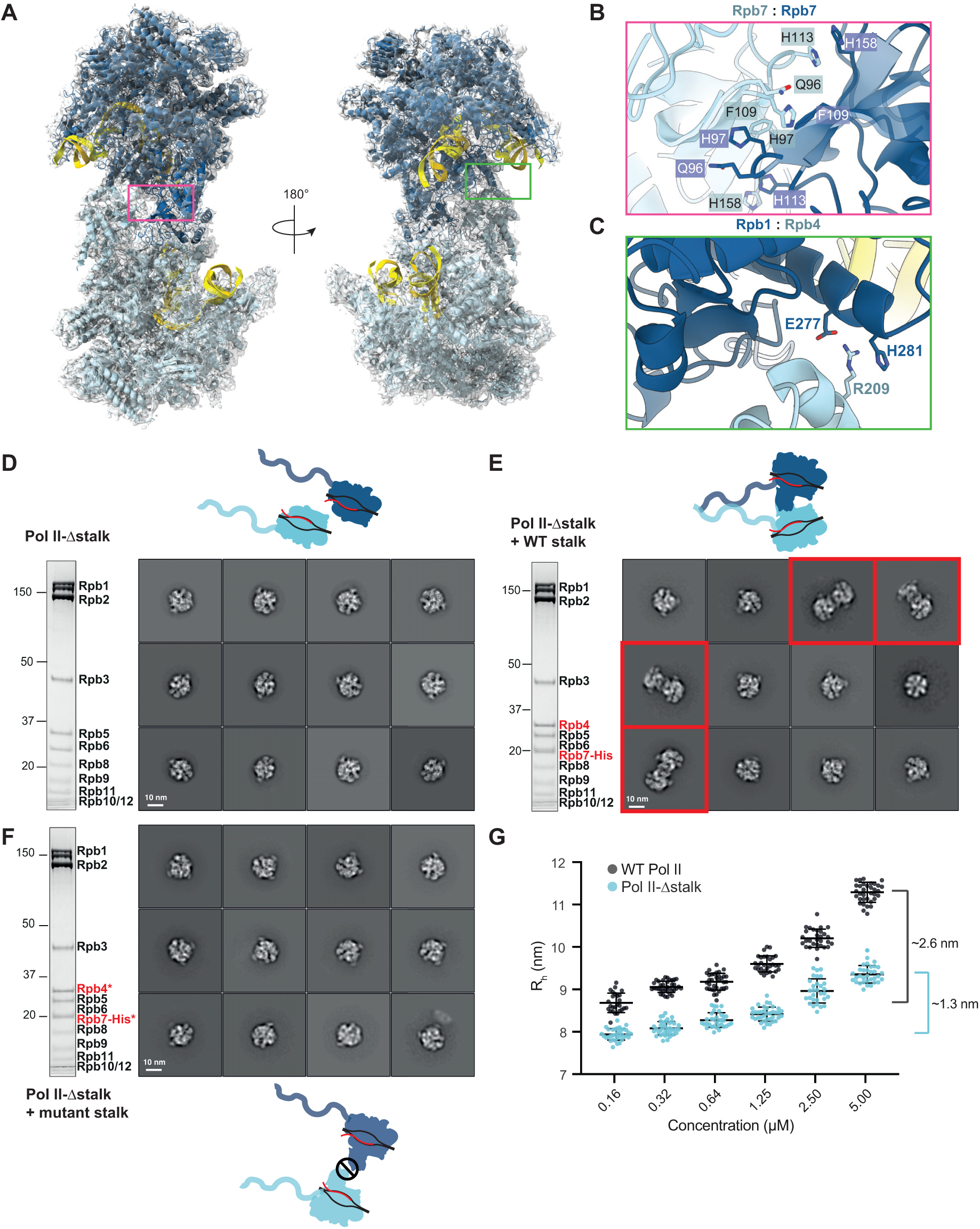
Pol II dimerizes via the stalk protein Rpb7. (**A**) Two copies of monomeric Pol II rigid body fit into the dimeric Pol II cryoEM density. Monomer 1, blue cartoon; monomer 2, cyan cartoon; DNA, yellow cartoon. The magenta and green boxes on the model indicate the close-up views of the Pol II dimer interfaces in panels **B** and **C** respectively. (**B**) Hydrophobic interaction between the Rpb7 subunits from each monomer. (**C**) Interaction between Rpb1 from monomer 1 (blue) with Rpb4 from monomer 2 (cyan). (**D-F**) SDS-PAGE and cryoEM analysis of Pol II dimerization. 2D class-averages were generated from 400,000 particles for Pol II- stalk (**D**), Pol II- stalk reconstituted with wild-type (WT) stalk (**E**), or Pol II- stalk complemented with the mutant stalk (**F**). Selected 2D classes are ordered by the number of particles per class, and dimeric classes are highlighted in red. On SDS-PAGE, recombinant Rpb4 and Rpb7-His are labelled in red and asterisks denote mutant stalk proteins. (**G**) Dynamic light scattering analysis on wild-type (dark grey) or stalk Pol II (cyan). The hydrodynamic radius (R_h_) is plotted as a function of Pol II concentration. Black bars represent mean ± s.d. on 40 acquisitions for each concentration point. The brackets on the right represent the increase in R_h_ from the lowest to the highest concentration for each condition. R_h_ values deriving from DLS measurements with weak scattering signal or showing multimodal distribution (on average 7 % per sample) were excluded from the plot.

The topology of the Pol II dimer is similar to a crystallographic dimer (Barnes *et al*., 2015) with the monomers rotated by ∼15° around the 2-fold symmetry axis (Movie 1). Pol II self associates via two interaction regions that together bury a surface area of ∼2,800 Å^2^ in the crystal structure. This includes a hydrophobic interaction between the Rpb7 stalk subunits of each monomer (Figure 3B), and a weaker polar interaction between the Rpb4 stalk subunit of one monomer and Rpb1 of the opposite monomer (Figure 3C). This stalk-to-stalk dimerization interface is adjacent to, but not overlapping with, the putative APT density in the crosslinked Pol II–APT complex (Figure S7B). Thus, crosslinking may stabilize APT binding but destabilize dimerization, and APT binding seems to be structurally compatible with Pol II dimerization. A Pol II dimer in which the cleft is occupied by the stalk of the opposite monomer was previously observed and was proposed to be an inactive, storage form of Pol II (Aibara et al., 2021). In contrast, the stalk-to-stalk dimer is structurally compatible with DNA–RNA loading.

To examine whether the interface in the crystallographic dimer is also involved in the dimerization of Pol II in solution, we introduced surface mutations in Rpb7 (Gln96Ala, His97Ala, Phe109Lys, His158Ala) and Rpb4 (Arg209Ala) (Figure 3B-C). As these mutations would likely not be tolerated *in vivo*, we first examined a recombinant version of wild-type and mutant Pol II stalk alone (Rpb4–Rpb7), expressed and purified from *E. coli.* Size exclusion chromatography coupled to multi angle light scattering (SEC-MALS) analysis is consistent with the wild-type stalk being in a monomer-dimer equilibrium. In contrast, the mutant stalk has an average molecular weight less than that of wild-type Rpb4–Rpb7, suggesting that mutations based on the crystallographic dimerization interface also destabilize the dimer in solution (Figure S7C).

To determine the effect of a disrupted stalk-to-stalk interface in the context of a fully reconstituted Pol II, we incubated DNA–RNA-loaded Pol II-Δstalk with recombinant Rpb4–Rpb7 (wild-type or mutant) and purified the resultant complex by gel filtration (Figure 3D-F). We then analyzed these Pol II variants by cryoEM and 2D classification. We did not observe any dimeric 2D classes with stalk-less Pol II or Pol II reconstituted with a mutant stalk (Figure 3D,F). However, Pol II dimerization was rescued after reconstitution with a wild-type stalk (Figure 3E, compare with Figure 2C).

We also performed dynamic light scattering to further examine Pol II behavior in solution. This showed that wild-type Pol II exhibits a concentration-dependent increase in hydrodynamic radius (R_h_) (Figure 3G), consistent with concentration-dependent formation of dimers in solution. This increase in R_h_ was reduced for Pol II-Δstalk suggesting that stalk deletion leads to defects in dimerization. Together, our analysis of Pol II indicates that transcribing Pol II can homodimerize through the stalk.

### CTD dephosphorylation promotes Pol II dimerization

Pol II dimerization is promoted by Ref2–Glc7–Swd2. However, since Ref2–Glc7–Swd2 was not visible in the cryoEM map of the Pol II dimer, it likely does not mediate dimerization by physically holding two Pol II monomers together. We therefore tested whether dephosphorylation of the Pol II CTD was required for dimerization.

First, we determined whether the CTD itself plays a role in dimerization. We performed cryoEM followed by 2D classification on Pol II-ΔCTD loaded with the DNA–RNA hybrid. This showed that there was a reduced frequency of Pol II-ΔCTD dimers compared to WT Pol II (Figure 4A, compare to Figure 2C). In agreement with this, Pol II-ΔCTD behaved similarly to the dimerization-defective Pol II-Δstalk in dynamic light scattering suggesting that deletion of the CTD causes defects in dimerization compared to WT Pol II (Figure 4B, compare to Figure 3G). Together, these data suggest that the presence of the CTD promotes Pol II dimerization.

**Figure 4.**
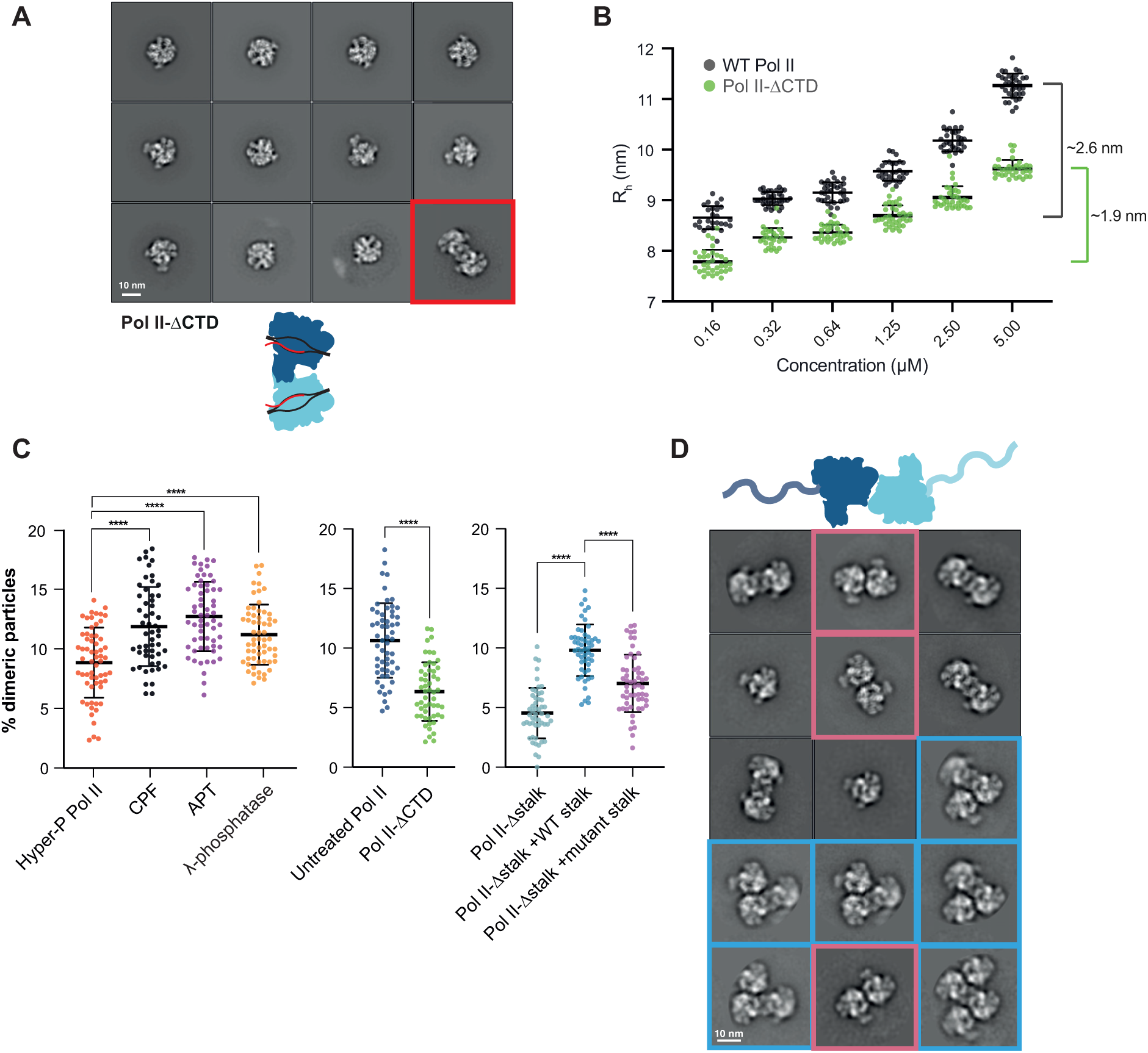
A dephosphorylated CTD promotes Pol II dimerization. (**A**) Selected 2D class-averages of Pol II- CTD from 400,000 particles, ordered by the number of particles in each class. A dimeric class is highlighted in red. (**B**) Dynamic light scattering analysis on wild-type (dark grey) or Pol II- CTD (green). Black bars represent mean ± s.d. on 40 acquisitions for each concentration point. The brackets on the right represent the increase in R_h_ between the lowest and the highest concentration for each condition. R_h_ values deriving from DLS measurements with weak scattering signal or showing multimodal distribution (on average 7 % per sample) were excluded from the plot. (**C**) Analysis of Pol II dimerization by negative stain EM. The percentage of Pol II dimeric particles over the total number of particles per micrograph is plotted. 60 micrographs were collected for each Pol II treatment (left). For the experiments in the middle and right panels 50 micrographs were acquired (see Figure S8A-B). All particles within each micrograph were identified by supervised Relion (v3.1) LoG-based autopicking (Zivanov et al., 2018). The experiments in the left and middle panels were performed twice, once as a ‘double-blind’ experiment. Black bars represent the mean ± s.d. Pairwise comparisons are shown as indicated where **** is P<0.0001 by one-way ANOVA Tukey’s test. n.s. = non-significant. (**D**) Selected cryoEM 2D-class averages of Pol II in the absence of a DNA–RNA hybrid. Classes with cleft-to-cleft Pol II dimers are highlighted in pink, whereas higher-order oligomers in light blue.

To determine whether the phosphorylation state of the CTD can modulate the extent of Pol II dimerization, we estimated the percentage of dimers under different conditions using negative stain EM. We hyperphosphorylated Pol II with Erk2 kinase and then incubated it with CPF, APT or λ-phosphatase (Figure S8A-B). We compared the percentage of dimers per micrograph in each condition, and calculated the mean dimerization frequencies to allow pairwise comparison (Figure 4C). This showed that CPF- and APT-treated Pol II had an increased frequency of dimerization compared to hyperphosphorylated Pol II. Treatment with λ-phosphatase also resulted in an increased number of dimers. Thus, Pol II dimerization is enhanced by phosphatase activity rather than the binding of CPF or APT.

We also calculated the proportion of dimers of Pol II-ΔCTD and Pol II-Δstalk and found that the dimerization frequency of both was decreased compared to wild-type, untreated Pol II (which is thought to be hypophosphorylated) (Figure 4C). In agreement with the cryoEM data, Pol II-Δstalk complemented with wild-type stalk has a higher proportion of dimers than Pol II with mutant stalk or Pol II-Δstalk in negative stain EM (Figure 4C right). Overall, these data show that dephosphorylated Pol II has an increased dimerization frequency compared to phosphorylated Pol II, and efficient dimerization requires the Pol II CTD and stalk.

Because negative stain EM is performed at low nanomolar concentrations, it was possible that some interactions were not retained during the experiments. We therefore analyzed Pol II in the absence of a nucleic acid scaffold as a control and found that it can form the stalk-to-stalk dimer, but it also dimerizes through a second interface (a cleft-to-cleft interaction) (Figure 4D). Since DNA occupies the cleft during transcription, the cleft-to-cleft interaction would not be possible on transcribing Pol II. In contrast, the stalk-to-stalk dimer is loaded on a DNA-RNA hybrid. Interestingly, we also observed higher-order oligomers which are formed by a combination of the stalk-to-stalk and cleft-to-cleft interactions (Figure 4D). Thus, Pol II can dimerize via at least two distinct mechanisms, and each of these may have different functional implications.

Taken together, our findings indicate that stalk-to-stalk Pol II dimerization is enhanced by CTD dephosphorylation, raising the interesting hypothesis that CTD phosphorylation state might regulate transcription by modulating Pol II dimerization.

### Ref2 binds Glc7 and acts as a PP1 regulator

Previous data suggested that Ref2 may regulate the Glc7 phosphatase within CPF (Nedea *et al*., 2008). PP1 phosphatases (including Glc7) function with a regulatory subunit that imparts substrate specificity (Peti et al., 2013). Regulatory subunits often interact with PP1 via an RVxF peptide motif. Additional interaction motifs are also commonly found in the regulatory subunits, for example the ΦΦ motif includes a pair of hydrophobic residues that bind PP1 (Fedoryshchak et al., 2020; Peti *et al*., 2013). The C-terminal region of Ref2 bears a non-canonical RVxF motif (SIKF in Ref2) that has been implicated in Glc7 binding *in vivo* (Nedea *et al*., 2008).

To examine the interaction between Ref2 and Glc7 *in vitro,* we generated C-terminal Ref2 truncations, and tested for co-migration with Glc7 in gel filtration (Figure S9A). The smallest truncation we tested comprised residues 348-405 of Ref2 (Ref2_348-405_) and we found that this interacted with Glc7. To understand the molecular basis of their association, we purified the Glc7–Ref2_348-405_ complex but this was refractory to crystallization. Therefore, we made a chimeric protein where the Ref2 fragment is fused to the flexible C-terminal tail of Glc7 (Figure 5A). We also included two repeats of the Pol II CTD in the chimeric protein with the aim of understanding how substrate is bound. We crystallized the chimeric protein and determined its structure at 1.85 Å resolution (Table 2, Figure 5B, Figure S9B). All residues of Glc7 and Ref2 could be modelled, except the last 14 residues of the flexible C-terminus of Glc7. The CTD substrate was not visible.

**Figure 5.**
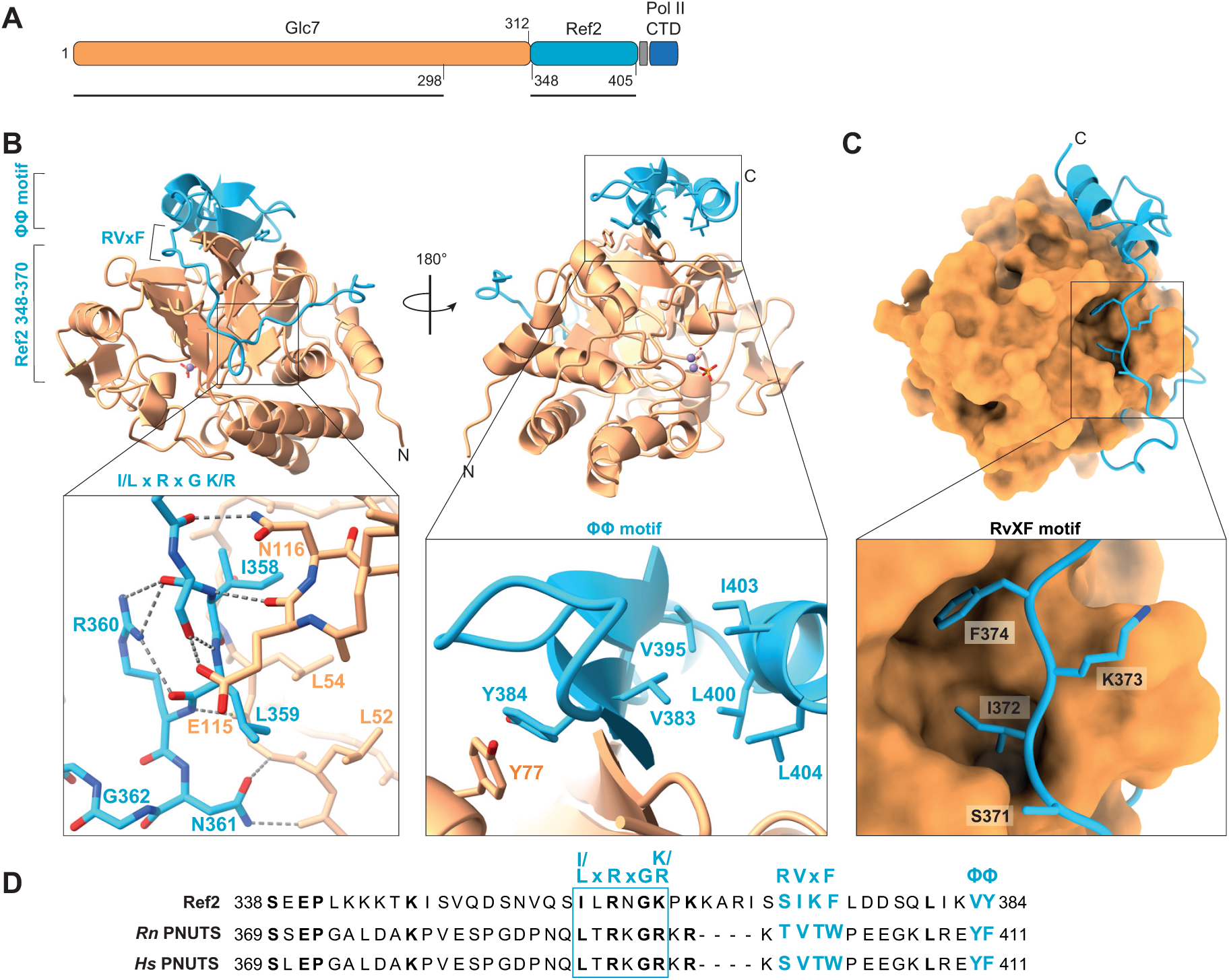
The Ref2 subunit of CPF and APT is a regulatory subunit of Glc7. (**A**) Schematic diagram of the Glc7–Ref2_348-405_ chimeric protein used for crystallization. Orange, Glc7; light blue, Ref2_348-405_; grey, glycine-serine linker; dark blue, two SPTYSPS Pol II CTD repeats. Black lines indicate the regions visible in the crystal structure. (**B**) Cartoon representation of the Glc7–Ref2_348-405_ crystal structure in two orientations, and close-up views of the interactions of Ref2 residues 358–362 (left) and of the intermolecular β-sheet formed between Glc7 and Ref2, including interaction details for the hydrophobic pair (ΦΦ) (right). Two manganese ions and a phosphate group are shown in ball-and-stick. Coordination waters are in red. The N- and C-termini are indicated. (**C**) View of the interaction interface between Ref2_348-405_ in cartoon, and Glc7 in surface representation. The inset shows how Ref2 binds Glc7 through the conserved ‘RVxF’ motif. The N- and C-termini are indicated. (**D**) Sequence alignment of Ref2 and putative orthologues from *Rattus norvegicus* (*Rn*) and *Homo sapiens* (*Hs*). The Ref2 ‘I/L-x-R-x-G-K/R’ motif is enclosed in a light blue box. The RVxF and downstream ΦΦ motifs shared among PP1-regulators are highlighted in light blue.

**Table 2.**
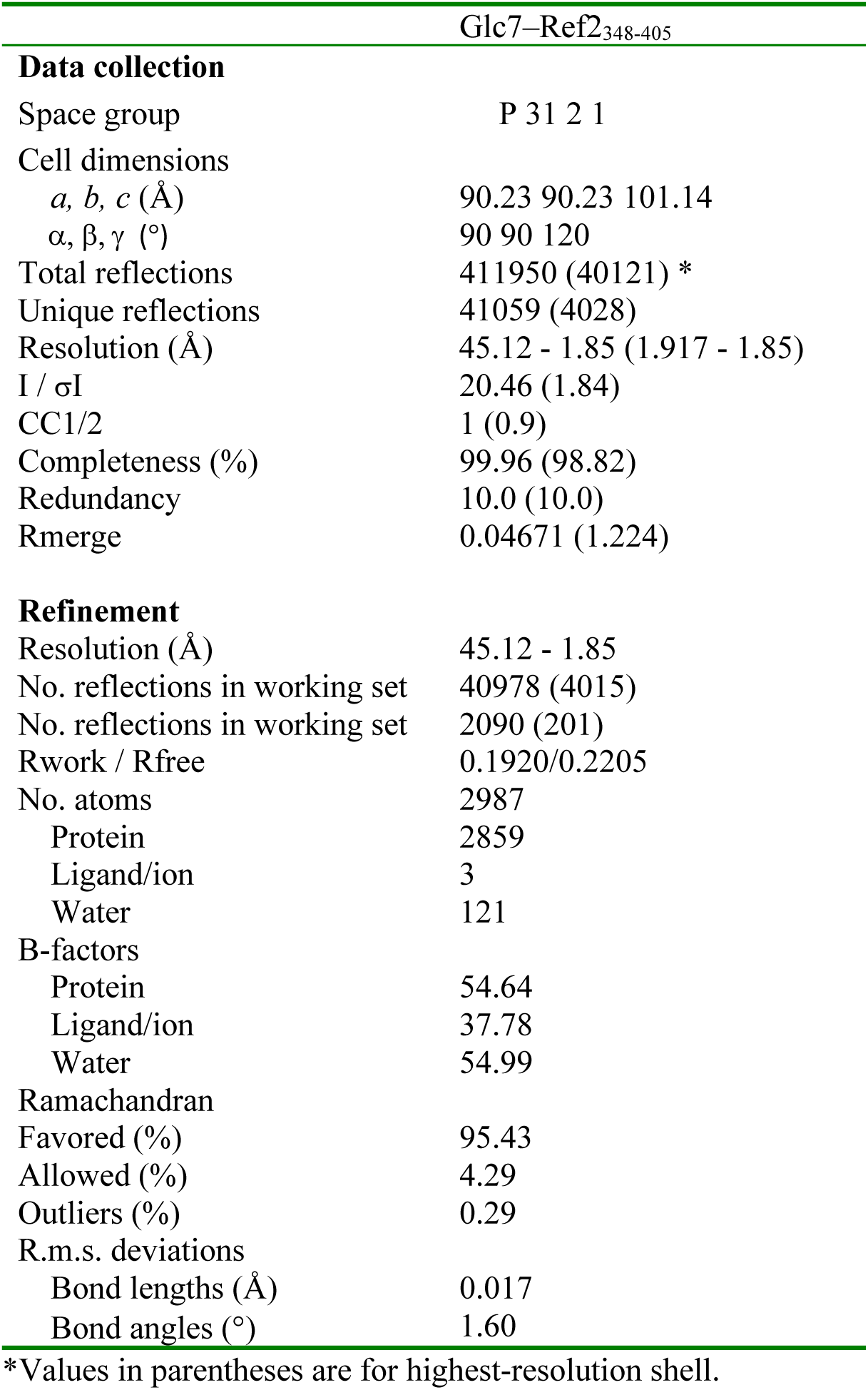
Data collection and refinement statistics (molecular replacement)

The structure shows that Ref2 wraps around the back of Glc7, burying a surface area of ∼4,200 Å^2^. The catalytic site on the opposite side of Glc7 is occupied by two manganese ions and one phosphate ion, as in the recently-reported structures of apo-Glc7 (PDB 7QWJ) and the Phactr1/PP1 holoenzyme (Fedoryshchak *et al*., 2020). The absence of density for the Pol II CTD in the active site could indicate weak binding affinity, crystal packing constraints, or a requirement for CTD modifications such as phosphorylation or proline isomerization (Werner-Allen et al., 2011; Xiang *et al*., 2010).

The extensive interaction between Ref2 and Glc7 encompasses three regions of Ref2 (Figure 5B-C). First, an N-terminal low complexity region (348-370) of Ref2 buries a surface area of ∼2,000 Å^2^ through polar and hydrophobic contacts with Glc7 (Figure 5B, left). Sequence alignment with putative Ref2 orthologues show that many of the interacting residues within this region are conserved (I/L-x-R-x-G-K/R) (Figure 5D). The other two Ref2–Glc7 interactions in the crystal structure are more generally conserved among PP1 regulatory subunits (Fedoryshchak *et al*., 2020; Peti *et al*., 2013). Within the non-canonical RVxF motif (SIKF in Ref2), Ile372 and Phe374 insert into a hydrophobic pocket on Glc7 (Figure 5C). This interaction is consistent with the previously-reported finding that Phe374 is required for Glc7 incorporation into the APT complex (Nedea *et al*., 2008). Finally, a pair of hydrophobic residues in Ref2, Val383 and Tyr384, form the ΦΦ motif that is found in many PP1 regulatory subunits (Figure 5B, right). Tyr384 stabilizes the formation of an intermolecular β-sheet via a π-π stacking interaction with Tyr77 of Glc7. Val383 inserts into a hydrophobic groove that includes residues 396-405 of Ref2, which form a C-terminal α-helix that folds back onto the Glc7–Ref2 β-sheet.

Superposition of the Glc7–Ref2_348-405_ crystal structure onto apo-Glc7 (Cα RMSD 0.47 Å) (PDB 7QWJ) or apo-PP1 (Cα RMSD 0.52 Å) (PDB 4MOV) indicates that Ref2 does not induce major conformational changes within Glc7 (Figure S9C). This is consistent with the mechanism of PP1 regulatory subunits, which often play roles in substrate recruitment, specificity and localization rather than as allosteric factors (Peti *et al*., 2013). In agreement with this, the *in vitro* dephosphorylation efficiency of hyperphosphorylated Pol II is similar for Glc7 and Glc7–Ref2_348-405_ (Figure S9D). Taken together, these findings suggest that Ref2 is a PP1 regulatory subunit.

### The Ref2 subunit of CPF and APT mediates the interaction with Pol II

We next aimed to determine whether Ref2 contributes to the interaction of CPF and APT with Pol II. Many Pol II interactions are mediated by intrinsically-disordered regions (IDRs) (Strzyz, 2018). Ref2 and its putative mammalian orthologue PNUTS (Choy et al., 2014; Cortazar *et al*., 2019; Shi et al., 2009) are predicted to be largely disordered by AlphaFold2, except for their TFIIS N-terminal domain (TND) which is a structured interaction module found in transcription elongation factors (Figure S10A) (Cermakova et al., 2021; Jumper et al., 2021; Varadi et al., 2022) . To test whether Ref2 is required for the interaction between APT and Pol II, we performed a pull-down assay using wild-type APT or APT lacking Ref2, and Pol II immobilized on StrepTactin beads. This revealed that the Ref2-containing version of APT, but not APT-ΔRef2, interacts with Pol II (Figure 6A). Thus, Ref2 is required for the interaction of APT (and presumably CPF) with Pol II.

**Figure 6.**
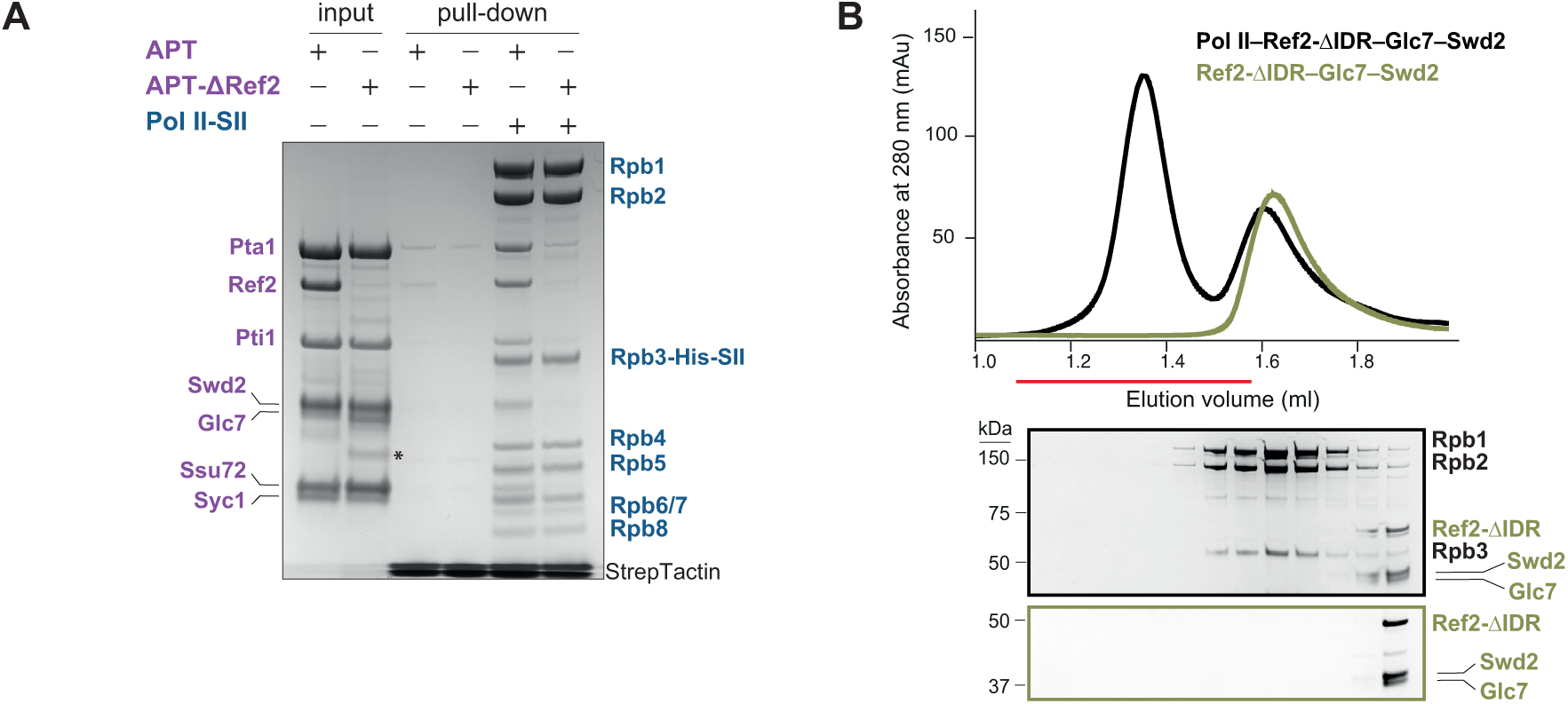
Ref2 is required for the APT interaction with Pol II. (**A**) Pulldown assay of APT and APT- Ref2 with Pol II immobilized on StrepTactin beads. SII indicates StrepII-tagged protein. APT subunits are labelled in purple and Pol II subunits are labelled in blue. APT- Ref2 was obtained after Ref2 was degraded by contaminating proteases during purification of APT. The asterisk indicates a degradation product. (**B**) Analytical size exclusion chromatography of Pol II and Ref2- IDR–Glc7–Swd2. The fractions indicated by a red line were analyzed on the SDS-PAGE below. Uncropped gels are in Supplementary Data 1B.

Next, we sought to determine which region of Ref2 is responsible for the interaction with Pol II. Given the emerging role of disordered regions in transcription regulation (Cermakova *et al*., 2021; Strzyz, 2018), we generated a Ref2-ΔIDR–Glc7–Swd2 complex where the IDR of Ref2 was removed. Size exclusion chromatography showed that the IDR of Ref2 is essential for the interaction of Ref2–Glc7–Swd2 with Pol II (Figure 6B). However, the interaction of APT with Pol II was not fully abrogated with Ref2-ΔIDR (Figure S10B), suggesting that APT might make multiple contacts with Pol II. Overall, these data show that Ref2 connects the Glc7 phosphatase subunit of CPF and APT to Pol II, thus creating a physical link between RNA 3ʹ-end processing and transcription regulation.

## Discussion

Here we describe physical and functional connections between the RNA 3ʹ-end processing and transcription machineries. We show that dephosphorylation of the Pol II CTD promotes stalk-to-stalk Pol II dimerization that is compatible with basal transcription. The Ref2 subunit of CPF and APT interacts with both Pol II and Glc7, acts as a PP1 regulatory subunit, and likely targets Glc7 activity to Pol II. Together, these findings provide new insight into the regulation and processing of eukaryotic RNAs.

The molecular role of Ref2 had been previously poorly characterized. In addition to a central IDR that mediates Pol II binding, and a C-terminal region that mediates Glc7/CPF/APT binding, Ref2 contains an N-terminal TND, a domain that mediates crosstalk between transcription factors (Cermakova *et al*., 2021) (Figure S10A). A region within the Ref2 IDR has also been reported to bind RNA (Russnak et al., 1996). Thus, Ref2 might act as a hub, bringing together the transcription machinery, the 3ʹ-end processing machinery, transcription factors and RNA. In humans and fission yeast, Ref2 and Glc7 orthologues also play roles in transcription termination (Cortazar *et al*., 2019; Kecman *et al*., 2018) and therefore the function of Ref2 is likely to be conserved.

CPF is thought to associate with Pol II and TFIID at promoters (Zhao *et al*., 1999) and remain associated throughout transcription (Lidschreiber *et al*., 2018). However, it had remained unclear if CPF interacts with Pol II directly. Other interactions between Pol II and the 3ʹ-end processing machinery have been previously reported. For example, within the CF IA complex, the Pcf11 subunit recognizes P-Ser2 of the CTD at the polyadenylation signal (Licatalosi *et al*., 2002; Pearson and Moore, 2014), and Pcf11–Clp1 binds the flap-loop of Rpb2 (Pearson and Moore, 2014). We show that, unlike CF IA and many transcription and RNA processing factors which are recruited by the CTD, CPF and APT contact core regions of Pol II, and the CTD is dispensable for their interaction.

We propose that CPF- and APT-mediated dephosphorylation of the Pol II CTD is highly regulated. CPF/APT may promote formation of the stalk-to-stalk dimer while Pol II is actively transcribing. Consistent with this, the stalk-to-stalk dimer forms on Pol II loaded onto a DNA–RNA transcription bubble. Association of APT near the RNA exit channel of Pol II (Figure 2A) would allow it to monitor the emerging RNA, as it is transcribed by Pol II (Figure 7A). The RNA capping enzyme complex (Martinez-Rucobo et al., 2015) and the U1 snRNP splicing complex (Zhang et al., 2021) also bind close to the RNA exit tunnel, but they all contact different parts of Pol II. Thus, our study shows that co-transcriptional RNA processing factors use a common mechanism of associating with transcribing Pol II near the RNA exit channel, but they achieve this through different contacts.

**Figure 7.**
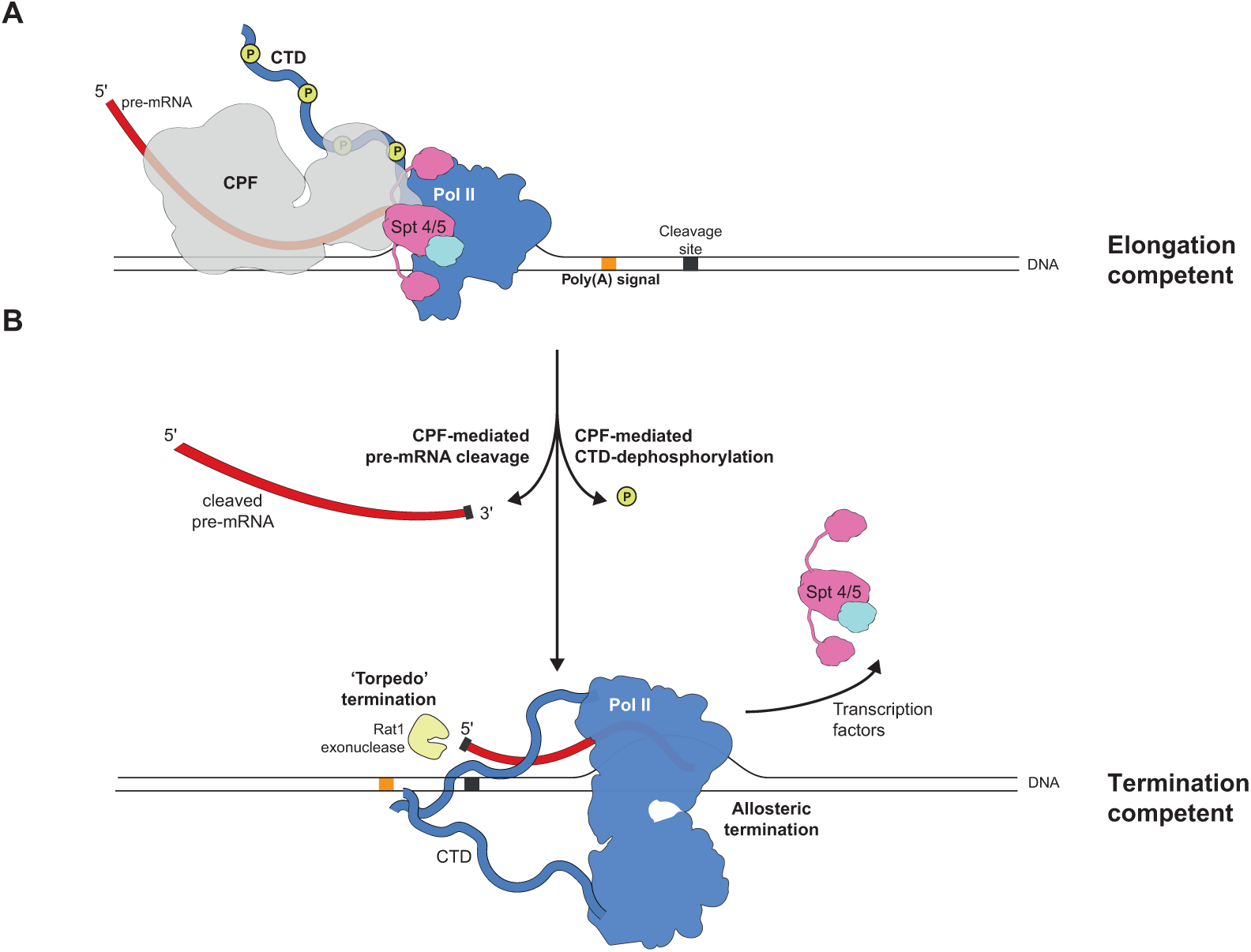
Proposed model for the role of CPF in transcription termination. (**A**) CPF and APT can bind elongating Pol II to monitor the nascent RNA as it emerges from the RNA exit channel. The poly(A) signal sequence (PAS) and the cleavage site are indicated by orange and grey boxes, respectively, on the template DNA. Nascent RNA is in dark red. Transcription elongation factors Spt4/5 are also shown. Phosphorylation marks are shown as yellows circles. (**B**) After the PAS is transcribed, CPF-mediated pre-mRNA cleavage and Pol II CTD-dephosphorylation are activated. As a result, the newly exposed 5’-end still attached to Pol II is degraded by the ‘torpedo’ exonuclease Rat1 (left), and Pol II dephosphorylation triggers an allosteric event (Pol II dimerization), which displaces transcription factors (right). Thus, CPF converts Pol II into a termination competent complex.

We propose that Pol II dimerization could be the structural change in Pol II that is postulated in the ‘allosteric’ model of transcription termination. The stalk-to-stalk Pol II dimer is not compatible with transcription initiation or elongation as there are major clashes with pre-initiation and elongation factors (Figure S11A-B). Thus, in a revised model of transcription termination, the 3ʹ-end processing machinery recognizes the cleavage site after transcription of the polyadenylation signal. This would activate Glc7 within CPF (or within APT on sn/snoRNAs) to dephosphorylate the Pol II CTD. Dephosphorylation promotes Pol II dimerization (the allosteric change), which is incompatible with elongation factor binding. Thus, this would render Pol II competent for transcription termination (Figure 7B).

After termination, multiple modes of Pol II dimerization (stalk-to-stalk and cleft-to-cleft) could contribute to the assembly of higher-order oligomers (Figure 4D), consistent with clustering of hypophosphorylated Pol II in biomolecular condensates (Boehning et al., 2018; Lu et al., 2018; Nosella and Forman-Kay, 2021; Rawat et al., 2021). In support of this possibility, DNA–RNA-loaded Pol II can undergo droplet-formation in the presence of APT *in vitro* (Figure S11D).

In summary, transcription of the RNA signals recognized by the 3ʹ-end processing machinery triggers a series of events that inhibit transcription elongation and promote transcription termination. A complex interplay between the nascent RNA, Pol II and the 3ʹ-end processing machinery suggests that regulation of transcription by RNA processing is likely underappreciated.

## Methods

### Cloning

#### Genome editing of S. cerevisiae RNA Polymerase II by CRISPR-Cas9

Endogenous yeast RNA Pol II from wild-type protease-deficient (JWY104) and Δ*RPB4* (OCSC2065) strains (Table S1), was purified by insertion of a His_6_-TEV-StrepII-tag (SII) at the C-terminus of Rpb3 via CRISPR-Cas9-mediated homologous recombination (Laughery and Wyrick, 2019). The same approach was used to introduce a 3C protease cleavage site within the *RPB1* ORF before the start of the CTD (residue 1454 of Rpb1) in JWY104. CRISPR-Cas9 engineering was performed based on Laughery and Wyrick (Laughery and Wyrick, 2019). First, we designed a 20 nt-long single guide RNA (sgRNA) matching the target DNA sequence harboring a protospacer-adjacent motif (PAM) within an 8-20 bp window from the site being edited. To design the optimal sgRNA for *S. pyogenes* Cas9 with a 5ʹ-NGG-3ʹ PAM site, we utilized the online resource CRISPOR (Concordet and Haeussler, 2018). Synthetic oligonucleotide sequences encoding the selected sgRNAs (Table S2) were cloned via *BclI* and *SwaI* restriction sites into a pML104 vector (Laughery et al., 2015), which expresses the Cas9 enzyme from *S. pyogenes* and carries a *URA3* marker. Double stranded DNA repair templates (donor DNA; Table S2) were designed with ∼30-40 bp homology arms and carried point mutations, preferably within the PAM, so that the target sequence will not be re-cleaved after the editing. 25 ng vector expressing sgRNA and 2 μg of the relevant donor DNA were co-transformed into yeast (Chen et al., 1992), and cells selected on SD/MSG agar -URA dropout plates. Colonies were screened for the intended modifications by PCR and sequencing. Positive clones were counter-selected on 5-fluoroorotic acid (5-FOA) containing plates to ensure removal the pML104 vector. All sgRNA and donor sequences are available in Table S2.

#### Generation of vectors carrying APT (sub)complexes for baculovirus

For APT, *E. coli* codon*-*optimized genes encoding Pta1 (Uniprot Q01329), Ref2-StrepII-tag (SII) (Uniprot P42073), Pti1 (Uniprot O49339), Swd2 (Uniprot P36104), Glc7 (Uniprot P32598), Ssu72 (Uniprot P53538), and Syc1 (Q08553) (Gene Art, Thermo Fisher) were amplified and cloned into pACEBac1 through *BamHI* and *XbaI* restriction sites. APT genes were subsequently amplified from pACEBac1 and assembled by Gibson assembly into vectors from a modified biGBac system (Hill et al., 2019; Weissmann et al., 2016). Syc1 and Pta1 were inserted into pBig1a; Ref2-SII, Pti1, Swd2, Glc7 and Ssu72 were inserted into pBig1b. Correct integration was verified by SwaI digestion. A pBig2ab vector containing all seven APT subunits was then assembled via a second Gibson reaction. The same approach was followed to clone the Ref2-SII, Glc7, Swd2 into pBig1a, and Pta1-SII, Pti1, Ssu72, Syc1 into pBig1b. For Ref2 IDR-SII–Glc7–Swd2, the IDR region of Ref2 (residues 230-338) was deleted.

#### Glc7 and Ref2 C-terminal truncations

Genes encoding Glc7 and Ref2_348-523_ were amplified from *E. coli* codon*-*optimized genes (Gene Art, Thermo Fisher) and cloned into FX cloning vectors carrying an N-terminal His_10_-SUMO tag (gift from Dr. Harvey MacMahon, MRC-LMB). Genes encoding Ref2_348-434_ and Ref2_348-405_ were amplified and cloned into the *BamHI* and *XhoI* sites of a pGEX-6P-2 vector, downstream of a GST-tag followed by a 3C cleavage site.

#### Glc7–Ref2_348-405_ fusion protein

The coding sequence for His_10_–SUMO–Glc7*–*Ref2_348-405_ followed by a Gly-Ser-Gly-Ser-Gly linker and two RNA Pol II CTD repeats (SPSYSPT) was synthesized and cloned into a pET24b expression vector by Epoch Life Science.

#### Recombinant Rpb4–Rpb7 (RNA Pol II stalk)

*RPB4* (Uniprot P20433) and *RPB7* (Uniprot P34087) genes were synthesized and cloned (Epoch Life Science) into a bicistronic pETDuet-1 vector. The gene encoding Rpb7-3C-His_6_ was inserted in the first cassette, and Rpb4 was inserted into the second. For the generation of the mutant Pol II stalk, point mutations were introduced by site-directed mutagenesis in *RPB4* (Arg209Ala) and in *RPB7* (Glu96Ala, His97Ala, Phe190Lys, His158Ala).

### Recombinant protein expression and purification from insect cells

Preparation of bacmids in EmBacY cells, transfection into Sf9 cells, initial and secondary baculovirus generation, and protein overexpression in Sf9 cells were carried out as previously described (Hill *et al*., 2019; Kumar et al., 2021). For large scale protein expression, 5-10 ml secondary virus was used to infect 500 ml suspension Sf9 cells (at 2×10^6^ cells/ml and >90% viability) grown in 2-litre roller bottle flasks. All Sf9 cells were grown in insect-EXPRESS (Lonza) media, incubated at 27°C and at 140 rpm. No additional supplements were provided to the media.

APT, the APT subcomplex Pta1-SII–Pti1–Ssu72–Syc1, CPF phosphatase module (Kumar *et al*., 2021), CPF-core (polymerase and nuclease modules) (Hill *et al*., 2019), the nuclease-phosphatase CPF module (Kumar *et al*., 2021) and CPF polymerase module (Casañal *et al*., 2017) were purified from Sf9 cell pellets according to the following protocol. Pellets from 4-litre cultures were harvested at 4,000 rpm in a JLA 8.1000 rotor and resuspended up to 120 ml with lysis buffer (50 mM Na-HEPES pH 8.0, 300 mM NaCl, 0.5 mM Mg(OAc)_2_, 0.5 mM TCEP and 10 % v/v glycerol) enriched with 4x EDTA-free protease inhibitor cocktail tablets (Roche, cat. No. 11836153001). Cells were lysed by sonication using a 10 mm tip on a VC 750 ultrasonic processor (Sonics). Sonication was performed at 70% amplitude with 5 seconds ‘on’ time and 10 seconds ‘off’ time and was followed by ultra-centrifugation at 45,000 rpm for 35 minutes in a Ti45 Beckman rotor. Clear lysates were treated with 4 ml BioLock (IBA, cat. No. 2-0205-050) before incubation with 2 ml bed volume of StrepTactin resin (IBA, cat. No. 2-1201-025) at 4 °C for 1 hour. Beads were washed with 100 ml lysis buffer, and proteins carrying a Strep-II tag were eluted by incubation with 20-25 ml lysis buffer supplemented with 6 mM desthiobiotin (IBA, cat. No. 2-1000-005). The filtered eluate (0.45 μM filter) was diluted to about 150 mM NaCl with buffer A (10 mM Na-HEPES pH 8.0, 0.5 mM TCEP and 5 % v/v glycerol) and loaded on a 6 ml Resource Q anion exchange chromatography column (Cytiva, cat. No. 17117901). Protein complexes were eluted by applying a gradient of 15–60 % buffer B (buffer A supplemented with 1M NaCl) over 7-8 CV. The relevant fractions were pooled, and proteins concentrated in Amicon Ultra-15 concentrators (Merck) before flash freezing and storage at - 80 °C.

For the APT subcomplex Ref2-SII–Glc7–Swd2 and Ref2 IDR-SII–Glc7–Swd2, the ion exchange chromatography step was replaced by gel filtration on a Superdex 200 Increase 10/300 GL (Cytiva, cat No. 28990944) equilibrated in 10 mM K-HEPES pH 8.0, 150 mM KCl, 0.5 mM TCEP and 5 % glycerol).

Recombinant CPF was reconstituted by incubating equimolar amounts of purified polymerase module and nuclease-phosphatase module on ice for 30 minutes, then injecting them on a Superose 6 Increase 10/300 GL column (Cytiva, cat No. 29091596) in 10 mM K-HEPES pH 8.0, 150 mM KCl, 0.5 mM TCEP and 5 % glycerol (Kumar *et al*., 2021). Peak fractions were analyzed by SDS-PAGE for proper stoichiometry, pooled, concentrated to 8.6 mg/ml, flash frozen and stored at -80 °C.

### Purification of endogenous RNA Polymerase II from *S. cerevisiae*

The purification of RNA Pol II from yeast was based on a previously described protocol (Sydow et al., 2009) with several modifications. For wild type (strain MCY102) and the Pol II -stalk (strain MCY101), 50-litre yeast cultures were grown in flasks in YEPD medium at 30°C, harvested at an OD_600_ of 8.0 (∼18-20 hours), resuspended in lysis buffer (200 mM Tris-HCl pH 8.0, 150 mM KCl, 10 μM Zn(OAc)_2_, 10 mM DTT, 1 % DMSO, and 10 w/v glycerol) supplemented with 5x protease EDTA-free inhibitor tablets (Roche, cat. No. 11836153001), and frozen by dropping the suspended yeast into liquid nitrogen. Lysis was performed through cryogenic grinding of the frozen cells by a Freezer Mill 6870 (SPEX CertiPrep). After thawing the ground material, the crude lysate was subjected to centrifugation for 30 minutes at 13,750 x g in a JLA 16.250 Beckman rotor. The supernatant was then ultracentrifuged at 45,000 rpm in a Ti45 Beckman rotor for 1.5 hours. (NH_4_)_2_SO_4_ was added to the cleared lysate until 50 % w/v saturation and precipitated overnight at 4 °C with gentle stirring. The precipitated solution was centrifuged at 33,768 x g in a JLA 16.250 Beckman rotor, and the supernatant was discarded. The (NH_4_)_2_SO_4_ pellet was resolubilized with a buffer without salt (50 mM Tris-HCl pH 8.0, 0.5 mM EDTA, 10 μM Zn(OAc)_2_, 7 mM imidazole, 2 mM DTT, and 10 w/v glycerol) supplemented with 1x protease inhibitor tablet, by adding 14 ml of buffer per 10 g pellet. The resuspended material was then subjected to centrifugation at 15,000 rpm in a JA 25.50 Beckman rotor for 10 minutes, passed through a 0.45 μM filter, and loaded on a 5 ml HisTrap HP column (Cytiva, cat. No. 17524802) for affinity purification. After washes, bound proteins were eluted with 200 mM imidazole over 30 CV, and the peak fractions were pooled for subsequent incubation with 1 ml (bed volume) pre-equilibrated StrepTactin resin (IBA, cat. No. 2-1201-025) for 1 hour and washed with buffer A containing 20 mM Tris-HCl pH 8.0, 0.5 mM EDTA, 150 mM KCl, 2 mM DTT, and 10 % glycerol. Pol II (Rpb3-SII) was eluted with 20 ml of buffer A supplemented with 8 mM desthiobiotin (IBA, cat. No. 2-1000-005), filtered through a 0.45 μM filter. It was then applied to a 1 ml MonoQ 5/50 GL column (Cytiva, cat. No. 17516601) and eluted with a linear gradient of 20–60 % buffer B (20 mM Tris-Acetate pH 8.0, 0.5 mM EDTA, 2 M KAc, 10 mM DTT, and 10 μM Zn(OAc)_2_, and 10 % glycerol) over 60 CV. Peak fractions were pooled and concentrated using a 30 kDa cut-off Amicon Ultra-15 concentrator (Merck, cat No. UFC903024) to a typical concentration of about 6 mg/ml, before flash freezing and storage at -80 °C.

For the Pol II Δ-stalk purification, the 50-litre culture was allowed to grow for ∼40 hours at 28°C and harvested at an OD_600_ of 5.0. Additional 5x protease EDTA-free inhibitor tablets (Roche, cat. No. 11836153001) and 5 µM Pepstatin A were added to the resuspension buffer to prevent CTD degradation.

For the yeast strain carrying a C-terminal His_6_-Bio tag on Rpb3 (BJ5464) the purification protocol was the same, except there was no incubation with StrepTactin resin (Sydow *et al*., 2009). After anion exchange, this version of Pol II required further polishing by gel filtration on a Superose 6 Increase 10/300 equilibrated in 10 mM K-HEPES pH 8.0, 100 mM KCl, 0.5 mM TCEP and 5 % glycerol.

### Recombinant protein expression and purification from *Escherichia coli*

#### His_10_–SUMO–Glc7 and His_10_–SUMO–Glc7–Ref2_348-405_–GSGSG–SPSYSPTSPSYSPT fusion protein

His_10_–SUMO–Glc7 and the Glc7*–*Ref2_348-405_ chimeric protein used for crystallography were purified from 36 litres of transformed Rosetta pLysS competent cells grown in 2xTY, induced with 0.25 mM IPTG at 18 °C for 18-20 hours, harvested, and resuspended in lysis buffer (50 mM Tris-HCl pH 8.0, 750 mM NaCl, 20 mM imidazole, 2 mM MnCl_2_, 1 mM TCEP 10 % v/v glycerol) supplemented with 4x EDTA-free protease inhibitor cocktail tablets (Roche, cat. No. 11836153001) and 4 μg/ml DNase I. After lysis by sonication and centrifugation at 45,000 rpm in a Beckman 45Ti rotor for 30 minutes, the clear lysate was applied to a 5 ml HisTrap HP column. Bound proteins were eluted with 300 mM imidazole after extensive washes with 40 mM imidazole. Fractions containing the His_10_–SUMO–Glc7*–*Ref2_348-405_–GSGSG–SPSYSPTSPSYSPT chimera were pooled and dialyzed overnight against 50 mM Tris-HCl pH 8.0, 500 mM NaCl, 2 mM MnCl**_2_**, 0.5 mM TCEP, 5% v/v glycerol in the presence of SUMO protease. The sample was concentrated down to 5 ml volume (Vivaspin, Sartorius) and further purified through a HiLoad 26/600 Superdex 200 pg (Cytiva) size-exclusion column equilibrated in the dialysis buffer. Peak fractions were analyzed by SDS-PAGE, pooled, concentrated to 13 mg/ml, and used for crystallization trials.

#### Recombinant Rpb4–Rpb7 expression and purification

Competent BL21(DE3) *E. coli* cells were transformed with 50 ng of the pETDuet vector carrying Rpb4 and Rpb7-3C-His_6_. For both wild type and mutant Rpb4–Rpb7, 6-litre cultures were grown in 2xTY medium to an OD of 0.8 and induced overnight at 18 °C with 0.25 mM IPTG. Pellets were resuspended up to 180 ml with lysis buffer (50 mM Tris-HCl pH 8.0, 300 mM NaCl, 20 mM imidazole, 0.5 mM TCEP, 10% glycerol) supplemented with 4x EDTA-free protease inhibitor cocktail tablets (Roche, cat. No. 11836153001) and 4 μg/ml DNase I, sonicated, and subjected to centrifugation at 45,000 rpm in a Ti45 Beckman rotor for 30 minutes. Cleared lysate was passed through a 0.45 μM filter, and loaded on a 5 ml HisTrap HP column, washed, and eluted with a buffer containing 20 mM Tris-HCl pH 8.0, 300 mM NaCl, 200 mM imidazole, 0.5 mM TCEP and 5% glycerol. Peak fractions were pooled, concentrated to 3 ml volume (Vivaspin, Sartorius), and further purified on a HiLoad 16/600 Superdex 200 pg (Cytiva) size-exclusion column equilibrated in 10 mM K-HEPES pH 8.0, 150 mM KCl, 0.5 mM TCEP, and 5% glycerol. Peak fractions were pooled and concentrated to 36 mg/ml prior to snap freezing and storage at -80 °C. To remove high-molecular weight contaminants, mutant Rpb4–Rpb7 was further polished on a 1 ml HiTRAP Heparin HP column (Cytiva, cat. No. 1704703) through a 7.5 to 50 % elution over 25 CV with a buffer consisting of 10 mM K-HEPES pH 8.0, 1M NaCl, 0.5 mM TCEP and 5 % glycerol. The protein complex was concentrated to 6.8 mg/ml and stored at -80 °C.

#### Ref2 C-terminal truncations expression and purification

Competent BL21(DE3) *E. coli* cells were transformed with 50 ng of the FX vector containing the sequence encoding His_10_-SUMO-Ref2_348-523_. 5-litre cultures were grown in 2xTY medium until an OD of 0.8 and induced overnight at 20 °C with 0.25 mM IPTG. Pellets were resuspended in 120 ml lysis buffer (50 mM Tris-HCl pH 8.0, 300 mM NaCl, 5 mM imidazole, 1 mM TCEP and 10 % glycerol) supplemented with 2x EDTA-free protease inhibitor cocktail tablets (Roche, cat. No. 11836153001) and 4 μg/ml DNase I, sonicated, and subjected to centrifugation at 45,000 rpm in a Ti45 Beckman rotor for 30 minutes. Cleared lysate was applied to 4 ml bed volume Ni-NTA Agarose beads (Qiagen, cat. No. 30210) for 1 hour. After washing beads with 20 mM Tris-HCl pH 8.0, 1 M NaCl, 20 mM imidazole, 1 mM TCEP and 5 % glycerol, His_10_-SUMO-Ref2_348-523_ was eluted by incubation with 40 ml buffer containing 20 mM Tris-HCl pH 8.0, 300 mM NaCl, 250 mM imidazole, 1 mM TCEP. After overnight cleavage with SUMO protease to remove the tag, the protein was diluted with buffer A (25 mM MES pH 6.5, 1 mM TCEP and 5 % glycerol) to about 50 mM NaCl for cation exchange chromatography. Ref2_348-523_ was then applied to a 1 ml Resource S column (Cytiva, cat. No. 17-1177-01), washed and eluted with a gradient of 5–40 % buffer B (25 mM MES pH 6.5, 1 M NaCl, 1 mM TCEP and 5 % glycerol) over 80 CV. Eluted fractions were pooled, concentrated to 2 ml volume in Amicon Ultra-15 concentrators (Merck), and injected on a HiLoad 16/600 Superdex 75 pg (Cytiva) size-exclusion column equilibrated in 25 mM MES pH 6.5, 200 mM NaCl, 1 mM TCEP and 5% glycerol. Purified protein fractions were pooled, concentrated, and de-salted for further purification on a 1 ml Heparin column (Cytiva). High-purity Ref2_348-523_ was eluted through a linear 4–50 % gradient of a buffer containing 1M NaCl, over 50 CV. Peak fractions were pooled, concentrated to 7 mg/ml and stored at -80 °C after snap freezing.

For Ref2_348-434_ anf Ref2_348-405_, 50 ng of the cloned pGEX-6P-2 were used to transform competent BL21(DE3) *E. coli* cells. 6-litre cultures were grown in 2xTY medium until an OD of 0.8 and induced overnight at 20 °C with 0.25 mM IPTG. Pellets were resuspended in 180 ml lysis buffer (50 mM Tris pH 8.0, 300 mM NaCl, 0.5 mM EDTA, 1 mM TCEP and 10 % glycerol), sonicated, and subjected to centrifugation at 45,000 rpm in a Ti45 Beckman rotor for 30 minutes. Cleared lysates were incubated with 5 ml bed volume Glutathione Sepharose 4B resin (GE Healthcare, cat. No. 17-0756-05) for 1 hour. Beads were washed with lysis buffer containing 1 M NaCl, de-salted in an 80 mM NaCl-containing buffer, and incubated overnight with 3C protease for GST-tag removal. Ref2_348-434_ or Ref2_348-405_ proteins were then loaded on a 1ml Resource S column, washed, and eluted with a gradient ranging from 8 to 30% buffer B (25 mM MES pH 6.5, 1M NaCl, 1 mM TCEP and 5 % glycerol) over 30 CV. Proteins were concentrated for injection on a HiLoad 16/600 Superdex 75 pg in 25 mM MES pH 6.5, 200 mM NaCl, 1 mM TCEP and 5% glycerol. Peak fractions were frozen and stored after concentration.

### *In vitro* Strep-Tactin pull-down assays

Reactions were set up in a total volume of 40 μl. 1.5 μM SII-tagged apo-RNA Pol II was immobilized on 10 μl (bed volume) of StrepTactin resin equilibrated in wash buffer (10 mM K-HEPES pH 8.0, 100 mM KCl, 0.5 mM TCEP, and 0.1 % v/v Tween-20). After rinsing to remove excess unbound bait, Pol II-bound beads were incubated with 3 μM of the indicated prey proteins for 1 hour on ice without shaking. Beads were then washed three times with 150 μl wash buffer (600 x g, 30 seconds, 4 °C), and bound species eluted by SDS-loading dye supplemented with 5% β-mercaptoethanol. Input for both bait and prey proteins were included in the gel as reference. The assays were resolved by SDS-PAGE (NuPAGE™ 4–12%, Bis-Tris, Invitrogen) run in MES-SDS buffer at 190 V for 50 minutes. For the pull-down experiment in Figure S1B, 1 μM SII-tagged CPF subcomplexes and APT were immobilized on beads, and incubated with 2 μM apo-Pol II in solution.

### Size exclusion chromatography (SEC) interaction studies

All analytical SEC binding assays were conducted on an ÄKTAmicro system (GE Healthcare) using a Superose 6 Increase 3.2/300 column (Cytiva, cat No. 29091598) equilibrated in 10 mM K-HEPES pH 8.0, 100 mM KCl, 0.5 mM TCEP, and 5% v/v glycerol. 2.5 μΜ apo-Pol II was incubated on ice for 10 minutes with a 1.8 molar excess of APT or APT-subcomplexes in a total volume of 50 μl and spun (5 minutes at 21,130 x g) before injection. The eluted volume was monitored by A_260-280_, collected in 50 μl fractions and analyzed by SDS-PAGE (NuPAGE™ 4 to 12%, Bis-Tris, Invitrogen) run in MES-SDS buffer at 190V for 50 minutes.

Interaction studies between Glc7 and Ref2 were carried out by incubating 5 μΜ Glc7 with 10 μΜ Ref2 C-terminal truncations in 50 μl. Co-migration was assessed after injection on a Superdex 200 Increase 3.2/300 column (Cytiva, cat No. 28990946). The eluted volume was collected in 50 μl fractions and analyzed on SDS-PAGE (Bolt™ 4 to 12%, Bis-Tris, Invitrogen) run in MES-SDS buffer at 200 V for 35 minutes.

### *In vitro* dephosphorylation assays on hyperphosphorylated Pol II

200 μg Pol II was hyperphosphorylated by incubation with 280 units of human MAP Kinase 2/Erk2 Protein (Merck, cat. No, 14-550) in a total volume of 50 μl in kinase buffer (50 mM Tris-HCl pH 7.4, 200 mM KCl, 5 mM MgCl_2_, 0.5 mM EGTA, 2 mM DTT) in the presence of 200 μM ATP at 30 °C for 60 minutes. Excess ATP and MAP kinase were removed by gel filtration on a Superose 6 Increase 3.2/300 column equilibrated in 10 mM K-HEPES pH 8.0, 100 mM KCl, 0.5 mM TCEP, and 5% v/v glycerol.

500 nM hyperphosphorylated Pol II was incubated with 1 μM CPF, APT, CPF/APT subcomplexes, Glc7 or Glc7–Ref2_348-405_ in 20 μl reactions in the presence of phosphatase buffer (20 mM K-HEPES pH 8.0, 150 mM KCl, 0.5 mM Mg(OAc)_2_, 0.5 mM MnCl_2_, 2 mM DTT). Reactions were incubated at 30 °C for 90 minutes and quenched by the addition of SDS-loading dye supplemented with 5 % β-mercaptoethanol. After heating at 70 °C, samples were analyzed by SDS-PAGE (Bolt™ 4 to 12%, Bis-Tris, Invitrogen), run in MES-SDS buffer at 200 V for 30 minutes, followed by immunoblot against phosphorylated Tyr1 or Ser5 CTD residues [antibodies 3D12 (1:7 in milk) and 3E8 (1:2000 in milk) respectively (Antibody Core Facility - Helmholtz-Zentrum Munich)]. Anti-Rpb3 [1Y26 antibody (abcam ab202893) 1:1000 in milk] was used as a loading control. Membranes in Figure S1A and Figure S8A (left) were incubated with fluorescent secondary antibodies (Goat anti-mouse IgG DyLight 800 ThermoFisher #SA5-10176 and Goat anti-rat IgG DyLight 800 ThermoFisher #SA5-10024), and images acquired on a LI-COR Odyssey instrument. Membranes in Figure S9D and Figure S8A (right) were incubated with HRP-conjugated secondary antibodies, developed by Amersham ECL Prime Western Blotting Detection Reagent (Cytiva, catalog # RPN2236), and images acquired on a ChemiDoc (Bio-Rad Laboratories).

### Crystallization and X-ray crystallography

The chimeric protein Glc7*–*Ref2_348-405_–GSGSG–2x(SPSYSPT) was diluted to 7 mg/ml in 10 mM Na-HEPES pH 8.0, 150 mM NaCl, 0.5 mM TCEP, 5% glycerol, and crystallization screens were set up using the MRC-LMB crystallization facility. Initial crystal hits were observed from experiments performed in 200 nl vapour diffusion sitting drops set up by a Mosquito Crystal nanodispenser (TTP Labtech) in MRC-2 well plates (Swissci, Hampton research). The best crystals grew in 40 % (v/v) 1,4-Butanediol and 0.1 M Tris-HCl pH 8.5, which could be reproduced manually in hanging drop consisting of 0.5 μl reservoir + 0.5 μl protein drop against 1 ml reservoir solution. Crystals were harvested and cryo-cooled without adding any cryoprotectant.

X-ray data were collected on beamline I04 at Diamond Light Source. 1800 images were recorded on a Dectris Eiger2 XE 16M detector using an oscillation range of 0.1° and an exposure time of 0.05 s per image (40% of beam transmission) at a temperature of 100 K. Datasets were initially processed with the Xia2 (Winter et al., 2013) automated pipeline, using XDS (Kabsch, 2010b) for indexing and integration, and XSCALE (Kabsch, 2010a) for scaling and merging. Data were cut at 1.85 Å resolution based on CC_1/2_ of 0.9 and I/σ of 1.84 in the highest resolution shell. The structure of Glc7*–*Ref2_348-405_–GSGSG–2x(SPSYSPT) was solved by molecular replacement in Phaser (McCoy, 2007) using apo-Glc7 (PDB 7QWJ) as a search model. The Ref2 fragment was interactively built *de novo* in COOT (Emsley et al., 2010) into the F_o_-F_c_ difference map, followed by iterative cycles of refinement in phenix.refine (Adams et al., 2010) and Refmac (Murshudov et al., 2011), and model adjustments in COOT. The final model was validated with MolProbity (Williams et al., 2018).

The buried surface area between Ref2 and Glc7 was calculated with the PDBePISA tool (Krissinel and Henrick, 2007). Images were prepared with UCSF ChimeraX (Pettersen et al., 2021) and PyMOL (The PyMOL Molecular Graphics System, Version 2.3.5, Schrödinger, LLC). Overlay of the Glc7*–*Ref2_348-405_ crystal structure to that of Glc7 and PP1, and calculation of their root-mean-square deviation (RMSD) of Cα were performed with PDBeFold (Krissinel & Henrick, 2004).

Protein sequence alignments of Ref2 orthologs in Figure 5D were conducted by Clustal Omega (RRID:SCR_001591). The structural features of the RVxF motif in human PNUTS and fission yeast PNUTS (Ppn1 or C74.02c, Uniprot: 074535) were predicted based on AlphaFold2 (Jumper *et al*., 2021; Varadi *et al*., 2022).

### Design of the DNA–RNA hybrid scaffolds

Nucleic acid scaffolds mimicking a transcription elongation bubble were designed for structural and functional studies based on sequences used in previous studies (Ehara *et al*., 2017) (Figure S2 and Table S2). All nucleotides were synthesized by Integrated DNA Technologies. For the transcription assays, a different DNA scaffold was used that annealed to the *CYC1d* sub-fragment of *CYC1* (Hill et al., 2019), and it encoded the downstream sequence of *CYC1* (Figure S2A). For the RNA cleavage assays, we modified the DNA scaffold to anneal to the 3’-UTR of the *in vitro* transcribed *CYC1* model pre-mRNA (Butler and Platt, 1988) (Figure S2B). For the EM studies on the Pol II–Ref2–Glc7–Swd2 complex, we used the scaffold reported in Ehara *et* al., 2017 (Figure S2C) with no further modifications. For the Pol II–APT EM studies, the scaffold from Ehara and colleagues was modified to anneal to the small-nucleolar RNA *snR47*, whose processing has been linked to APT (Lidschreiber *et al*., 2018) (Figure S2C). All the template strand (TS) and the non-template strand (NTS) DNAs were designed to have an 11 bp mismatch, to allow the formation of a stable 9 nt DNA:RNA hybrid in the transcription bubble (Kireeva et al., 2000).

The RNA was first annealed to TS-DNA at 40 μM concentration in 20 mM Tris-HCl pH 7.5, 4 mM Mg(OAc)_2_, and 150 mM KAc in 1.5 ml Eppendorf amber tubes to protect the fluorophore. The reaction was heated to 95 °C for 5 minutes, and then allowed to gradually cool at room temperature. The TS-DNA–RNA scaffold was subsequently mounted step-wise on hyperphosphorylated Pol II (Kireeva *et al*., 2000). Pol II was incubated with 1.5 molar excess of the TS-DNA–RNA hybrid in 10 mM K-HEPES pH 8.0, 100 mM KCl and 0.5 mM TCEP for 10 minutes at 30 °C with gentle shaking (300 rpm). An equal amount of NTS-DNA strand was added to the reaction, and the 10-minute incubation was repeated. After centrifugation for 5 minutes at 21,130 x g, the sample was subjected to gel filtration in 10 mM K-HEPES pH 8.0, 100 mM KCl, 0.5 mM TCEP and 5 % glycerol to remove the unbound nucleic acids. Peak fractions with an A_260/280nm_ ∼1 were pooled, concentrated and stored as DNA–RNA-loaded Pol II to be utilized in EM (see EM methods).

### Pol II *in vitro* transcription assay

For the RNA extension assays in the presence of APT and CPF we employed the DNA-RNA hybrid displayed in Figure S2A and Table S2. Hyperphosphorylated Pol II was assembled with the scaffold as detailed above and subjected to gel filtration. The DNA–RNA-loaded Pol II was then incubated with a 0.5 mM mixture of nucleotide triphosphates (NTPs) in 50 μl reactions in transcription buffer (20 mM Tris-HCl pH 8.0, 150 mM KAc, 5 mM Mg(OAc)_2_. 5 μl fractions were taken over time to monitor the progression of transcription, and quenched in a stop solution (130 mM EDTA, 5% (w/v) SDS, and 12 mg/ml proteinase K) at 37 °C for 10 minutes. 2 x denaturing loading dye (95% formamide, 10 mM EDTA, 0.01% w/v bromophenol blue) was added to the samples before loading onto 20% TBE (Tris-borate-EDTA)-polyacrylamide gels containing 7 M urea. The gels were run at 400 V in 1 x TBE running buffer for 45 minutes and were scanned with an Amersham™ Typhoon™ Biomolecular Imager for 5′ 6-FAM labelled *CYC1* RNA with excitation at 488 nm.

### RNA *in vitro* cleavage assay

CPF cleavage assays were performed at 30 °C in a reaction buffer containing 10 mM K-HEPES pH 8.0, 100 mM KCl and 0.5 mM TCEP on 100 nM of the 259-nt *CYC1* pre-mRNA annealed to an artificial transcription bubble as described above (Figure S2B), in the absence or in the presence of Pol II. Purified recombinant CPF was added to a final concentration of 150 nM in the reaction. The cleavage factors, CF IA and CF IB were diluted to 15 µM in 20 mM Na-HEPES pH 8.0, 250 mM NaCl, and 0.5 mM TCEP and added to a final concentration of 450 nM in the reaction. Reactions were stopped at different time points as indicated, by incubation for 10 minutes at 37 °C in a stop solution containing 130 mM EDTA, 5% (w/v) SDS, and 12 mg/ml proteinase K in reaction buffer. 2 x denaturing loading dye (95% formamide, 10 mM EDTA, 0.01% w/v bromophenol blue) was added to the samples before loading onto 15% TBE (Tris-borate-EDTA)-polyacrylamide gels containing 7 M urea. The gels were run at 400 V in 1 x TBE running buffer, stained with SYBR Green (Invitrogen S7564), and imaged using an Amersham™ Typhoon™ Biomolecular Imager.

### Negative Stain Electron Microscopy

#### Specimen preparation

To compare the extent of Pol II dimerization under different conditions we used negative stain electron microscopy. 200 μg Pol DNA–RNA-loaded Pol II (Ehara *et al*., 2017 and Figure S2C) was hyperphosphorylated and subjected to SEC in 10 mM K-HEPES pH 8.0, 100 mM KCl and 0.5 mM TCEP as reported above. After concentration, 50 μg hyperphosphorylated Pol II was incubated with with 1 μM of either CPF or APT in a total volume of 50 μl for 90 minutes at 30 °C in phosphatase buffer (20 mM K-HEPES pH 8.0, 150 mM KCl, 0.5 mM Mg(OAc)_2_, 0.5 mM MnCl_2_, 2 mM DTT). For the same set of experiments, 50 μg hyperphosphorylated Pol II was was dephosphorylated with 250 units of λ-phosphatase (NEB, cat. No. P0753L) supplemented with PMP buffer and MnCl2 in a total 50 μl volume for 30 minutes at 30 °C. Pol II-ΔCTD was generated by incubation of unphosphorylated Pol II with 2.5 μg 3C protease in 50 μl. Purified Pol II Δ-stalk was incubated with 3 molar excess of recombinant Rpb4–Rpb7 (wild type or mutant). After treatment, each Pol II-containing reaction was loaded on gel filtration (Superose 6 3.2/300 equilibrated in 10 mM K-HEPES pH 8.0, 100 mM KCl and 0.5 mM TCEP) in order to remove excess reagents, monitor Pol II integrity, and apply the same buffer to all conditions being compared. Peak fractions were pooled, and Pol II diluted to 20 nM in 10 mM K-HEPES pH 8.0, 100 mM KCl, 0.5 mM TCEP and 0.005 % v/v Tween-20) for negative stain specimen preparation. 3 μl sample was applied to copper 400-mesh grids with an ultra-thin layer of carbon (Electron Microscopy Sciences, cat. No. CF400-Cu-UL), that had been glow-discharged in air for 30 seconds. The sample was incubated on the support for 60 seconds and then wicked with filter paper. Grids were sequentially applied to three 30 μl drops of 2% w/v uranyl acetate (7 seconds floating on top of each drop), and the excess stain removed by filter paper.

#### Data collection and processing

Grids were stored until imaging on a Tecnai Spirit microscope (FEI) operated at 120 keV, equipped with an Orius SC200W CCD camera (Gatan). 50-60 micrographs for each Pol II treatment were collected at a magnification of 21,000 X (2.53 Å/pixel), at -1 μm defocus and a total electron dose of 40-60 e^-^/A^2^ over 2 seconds. For each micrograph, total particle auto-picking was performed with Laplacian-of-Gaussian (LoG) in Relion (v3.1), using a 300 Å-wide area, which can accommodate Pol II dimers (230 Å through the longest axis). Automated picking was manually inspected to check the quality of the particles considered as ‘dimeric, using the 2D-projections of the cryoEM Pol II dimer as a reference (Figure S8C). Trimeric particles, if present, were also picked as dimers. For each Pol II sample, the ratio of dimeric particles over the total number of particles per micrograph was determined and plotted to calculate means and standard deviations using Graphpad Prism 9. Means were used for pairwise comparison between treatments, and statistical significance assessed by one-way ANOVA Tukey’s test. The quantifications in the left and middle panels of Figure 4 were performed twice in ‘double-blind’ replicates.

### Single-Particle Electron Cryo-Microscopy

#### Specimen preparation and data collection

**Pol II–APT** - To stabilize the DNA–RNA-loaded Pol II (scaffold in Figure S2C) in the presence of APT, we utilized gradient centrifugation coupled to cross-linking (GraFix) (Kastner *et al*., 2008; Stark, 2010). A 10–30 % sucrose gradient in the presence of 0.1 % glutaraldehyde (Figure S3A) was generated with a Biocomp Gradient Station. 200 ng sample in 100 μl was pipetted on top of the gradient before ultracentrifugation at 33,000 rpm in a SW40 Ti Beckman rotor at 4 °C for 18 hours. The sample was then manually fractionated, quenched in 100 mM Tris-HCl pH 8.0 and analyzed on a NuPAGE™ 3–8 % Tris-Acetate gel (Invitrogen). Fractions containing protein bands corresponding to the expected size of a Pol II–APT complex were dialyzed against 10 mM K-HEPES pH 8.0, 100 mM KCl and 0.5 mM TCEP (final concentration of 200–300 nM).

**Pol II–Ref2–Glc7–Swd2** - 2.5 μM DNA–RNA-loaded Pol II (Ehara *et al*., 2017 and Figure S2C) was assembled with 1.8 molar excess of Ref2–Glc7–Swd2 and incubated on ice for 30 minutes in 50 μl. After spinning at 21,130 x g, the supernatant was injected on a Superose 6 3.2/300 in 10 mM K-HEPES pH 8.0, 100 mM KCl and 0.5 mM TCEP. Pooled fractions were concentrated in Vivaspin 500 (Sartorious, cat. No. VS0121) to 2 μM. 0.005 % v/v Tween-20 was added just before vitrification.

**Pol II Δ-stalk and Pol II Δ-CTD** was prepared in the same conditions detailed above in the ‘negative stain EM’ session, and concentrated to 2 μM. 0.005 % v/v Tween-20 was added just before vitrification.

For all protein complexes and Pol II deletions, 3 μl sample was applied to 1.2/1.3 or 0.6/1 UltrAuFoil grids (Quantifoil) (Russo and Passmore, 2014) made hydrophilic using a 9:1 Argon:Oxygen plasma for 45 seconds. The applied sample was blotted for 5 seconds with a blot force of -12 at 4 °C and 100 % humidity, and plunged into liquid ethane on a Vitrobot Mark IV (FEI). Grids were imaged on Titan Krios microscopes operated at 300 keV using the EPU automated data collection software. Total dose (40 e^-^/Å^2^) and defocus range (−1.5 μm to -2.7) were kept consistent throughout datasets and microscopes.

For **Pol I–APT**, 7,980 movies were collected on a Gatan K3 detector in super-resolution mode, with a pixel size of 0.83 Å/pixel on Krios III at eBIC.

For **Pol II–Ref2–Glc7–Swd2**, a total of 27,500 movies were recorded. Initial datasets with stage tilt (0 °, 30 ° and 40°) were collected on Krios I (MRC-LMB) with a Falcon III detector in counting mode, with a pixel size of 1.04 Å/pixel. Data at zero tilt were collected at the University of Cambridge with a Gatan K3 detector in super-resolution mode, with a pixel size of 0.83 Å/pixel. Data with a tilted stage (33°, 40 ° and 45°) were acquired at MRC-LMB on Krios III with a Gatan K3 detector in counting mode, pixel size of 0.86 Å/pixel.

For **Pol II Δ-stalk and Pol II Δ-CTD**, 1,200 movies for each condition (containing ∼400,000 particles) were collected on Krios I (MRC-LMB) with a Falcon III detector in integrating mode, with a pixel size of 1.04 Å/pixel.

#### Data processing

For all samples, the stack of frames from the movies were corrected for whole-frame motion using Relion’s version of MotionCor2 (Zheng et al., 2017), and the defocus was estimated using CTFFIND4 (Rohou and Grigorieff, 2015). A detailed processing flowchart for the **Pol II–APT** is shown in Figure S3C.

For **Pol II–Ref2–Glc7–Swd2**, all seven collected datasets were treated separately for initial processing. The best particles were merged only after 2D classification, keeping Falcon III and K3 data separate, as described in the processing schematic (Figure S5). Image processing was performed in Relion (v3.1) (Zivanov *et al*., 2018) and Relion wrappers were used for external programs except for crYOLO. Particles were picked using a general network model in crYOLO (Wagner et al., 2019). For each dataset, three rounds of 2D classification on 3.5x binned particles were performed to remove junk particles. High-resolution classes containing both Pol II monomers and dimers were then merged and used to generate an initial 3D model. The initial model was used as a reference for one round of 3D classification to separate monomers from dimers. The Pol II dimer classes were subjected to 3D refinement with no symmetry applied. Particles were unbinned and the Falcon III and K3 datasets were merged with a consensus pixel size of 0.83 Å^2^/pixel. Merged particles were refined further resulting in a 4.6 Å 3D reconstruction of the Pol II dimer.

Focused refinement on each monomer was performed after signal subtraction from the other monomer. The two separately focus-refined monomers showed no noticeable difference between each other. Therefore, to further improve the resolution of our map, we applied ‘local symmetry’ (Shakeel et al., 2019), whereby particles from each monomeric subunit of the dimer were merged after signal subtraction of the other subunit. We refined this merged particle stack against the ‘focus-refined’ map of either monomer yielding 3.6 Å maps irrespective of which monomer was used as the reference. Maps were post-processed using negative B-factor values calculated by Relion. Map validation according to the gold-standard FSC at 0.143, and local resolutions maps are displayed in Figure S6.

Two copies of a monomeric Pol II crystal structure (Barnes *et al*., 2015) were manually fitted into the EM density of the Pol II dimer and then subjected to rigid fitting in USCF Chimera (Pettersen et al., 2004). Illustrations and movies were prepared using UCSF ChimeraX (Pettersen *et al*., 2021) and PyMOL (The PyMOL Molecular Graphics System, Version 2.3.5, Schrödinger, LLC). The buried surface area between the Rpb7 subunits within the Pol II crystallographic dimer (PDB 5C4X) was calculated in PyMOL with the ‘get_area’ command.

### Dynamic Light Scattering (DLS) analysis

To analyze the concentration-dependent dimerization of Pol II, Pol II-Δstalk and Pol II-ΔCTD were analysed by DLS. DNA–RNA hybrid-loaded Pol II (Ehara *et al*., 2017 and Figure S2C) was serially diluted from 5 μM to 156 nM in 10 mM K-HEPES pH 8.0, 100 mM KCl and 0,5 mM TCEP. 25 μl of each dilution was applied to a 384-well black/clear bottom Polystyrene microplate (Corning, cat No. 3540), followed by centrifugation to remove air bubbles or aggregates in the samples (2 minutes, 4,000 x g). 40 consecutive DLS measurements were acquired for each concentration point on a DynaPro plate reader (Wyatt Technology) at 25 °C. Data were analyzed directly in the accompanying DYNAMICS® software, and an autocorrelation function calculated for each measurement. The translational diffusion coefficient was obtained through automated nonlinear least squares fitting of the autocorrelation function, and the hydrodynamic radius (R_h_) derived through the Stokes-Einstein equation. Visualization of the R_h_ as a function of protein concentration was carried out in GraphPad Prism version 9.0. R_h_ values deriving from DLS measurements with weak scattering signal or showing multimodal distribution (on average 7 % per sample) were excluded from the plots.

### SEC Multiangle Light Scattering (MALS) analysis

The mass in solution of both wild type and mutant Rpb4–Rpb7 (RNA Pol II stalk) was determined by SEC-MALS measurements using a Wyatt Heleos II 18 angle light scattering instrument coupled to a Wyatt Optilab rEX online refractive index detector. Protein samples (100 μl at 2 mg/ml) were resolved using a Superdex 200 10/300 analytical gel filtration column (GE Healthcare) running at 0.5 ml/min at 25 °C in 10 mM K-HEPES pH 8.0, 100 mM KCl and 0.5 mM TCEP. Protein concentration was determined from the excess differential refractive index (RI) based on 0.186 RI increment for 1 g/ml protein solution. The concentration and the observed scattered intensity at each point in the chromatograms were used to calculate the absolute molecular mass from the intercept of the Debye plot using the Zimm model as implemented in Wyatt’s ASTRA software.

### *In vitro* phase separation and microscopy

Droplet formation occurs upon mixing 0.6 μM DNA–RNA-loaded Pol II with ≥ 4 μM APT (titration experiment, not shown). To follow the distribution of Pol II and RNA in the condensates, in the presence or in the absence of APT, we used confocal microscopy. The RNA annealed to the nucleic acid scaffold was labelled at the 5ʹ with fluorescein (5′ 6-FAM) (Integrated DNA Technologies). For Pol II, we took advantage of the Rpb3-His_6_-Bio tag carrying a biotinylation acceptor peptide, which can be biotinylated *in vitro* by incubation with bacterial BirA biotin ligase (Merck, cat. No. SRP0717). Biotinylation enabled us to label Pol II via Streptavidin-conjugated Alexa Fluor^TM^ 647 (ThermoFisher, cat. No. S21374). The droplets were allowed to form for 2 minutes in a 0.5 ml Protein LoBind Eppendorf tube in 10 mM K-HEPES pH 8.0, 100 mM KCl and 0,5 mM TCEP. 4 μl of the reaction was transferred to buffer-rinsed 50-well CultureWell chambered coverslides (Grace Bio-Labs, cat. No. GBL103350), and sealed with crystal clear tape. Imaging was conducted on a Zeiss LSM 780 laser scanning confocal microscope with a Plan-Apochromat 20x/0.8 M27 objective, by focusing on the bottom of the coverslide, where droplets settle. The sample was imaged with an excitation wavelength of 488 nm and 633 nm for FAM-RNA and Pol II-Alexa Fluor 647, respectively. Images were further processed and merged in FIJI (version 2.0.0-rc-66/1.52b).

## Acknowledgements

We thank S. Shakeel, S. Scheres, A. Kumar, K. Malhotra, G. Lee, T. Tang as well as P. Alcón, J. Stowell, J. Rodriguez Molina, and all the other members of the Passmore lab for assistance and advice; J.G. Shi for baculovirus; J. Grimmett and T. Darling (LMB scientific computation) for support; J. Howe (LMB light microscopy) for guidance with confocal microscopy. We acknowledge Olga Calvo and Patrick Cramer for sharing yeast strains, and Core Facility Monoclonal Antibodies at Helmholtz Munich for providing the 3D12 antibody. We acknowledge the MRC Laboratory of Molecular Biology Electron Microscopy Facility for access and support of electron microscopy sample preparation and data collection; and K. Cunnea, Y. Song and Diamond Light Source for access to eBIC (BI23268) funded by the Wellcome Trust, MRC and Biotechnology and Biological Sciences Research Council. We acknowledge D. Chirgadze and S.W. Hardwick (Cryo-EM facility, University of Cambridge) for cryoEM data collection. We thank D. Barford, H. Mischo, A. Tufegdžić Vidaković and S. West for careful reading of the manuscript and feedback. This work was supported by the Medical Research Council, as part of United Kingdom Research and Innovation, MRC file reference number MC_U105192715 (L.A.P.); the European Union’s Horizon 2020 research and innovation programme (ERC Consolidator grant agreement 725685) (to L.A.P); an EMBO long-term fellowship to M.C. (EMBO ALTF 556-2018); and a Novo Nordisk Fonden grant (NNF19OC0054219) to M.C.M.

## Author contributions

M.C. designed and performed experiments, analyzed the data and wrote the manuscript. M.C.M. conducted the in vitro RNA cleavage assays. M.C. and D.B. collected X-ray diffraction data and determined the structure of Ref2–Glc7. L.A.P. supervised the project, analyzed the data and wrote the manuscript.

## Competing interests

L.A.P. is an inventor of a patent filed by the Medical Research Council for all-gold EM support, licensed to Quantifoil under the trademark UltrAuFoil.

## Data availability

CryoEM maps of the dimeric yeast RNA polymerase II bound to a transcription bubble were deposited in the Electron Microscopy Data Base (EMDB) with the following accession codes: EMD-15358 (consensus map), EMD-15359 (focused map for monomer 1), and EMD-15360 (focused map for monomer 2). Atomic coordinates and structure factors for the Glc7–Ref2_348-405_ crystal structure were deposited in the Protein Data Bank (PDB) with the accession code PDB 8A8F. Correspondence and requests for materials should be addressed to L.A.P. All unique materials are available upon request with completion of a standard Materials Transfer Agreement.

---

**Figure S1.**
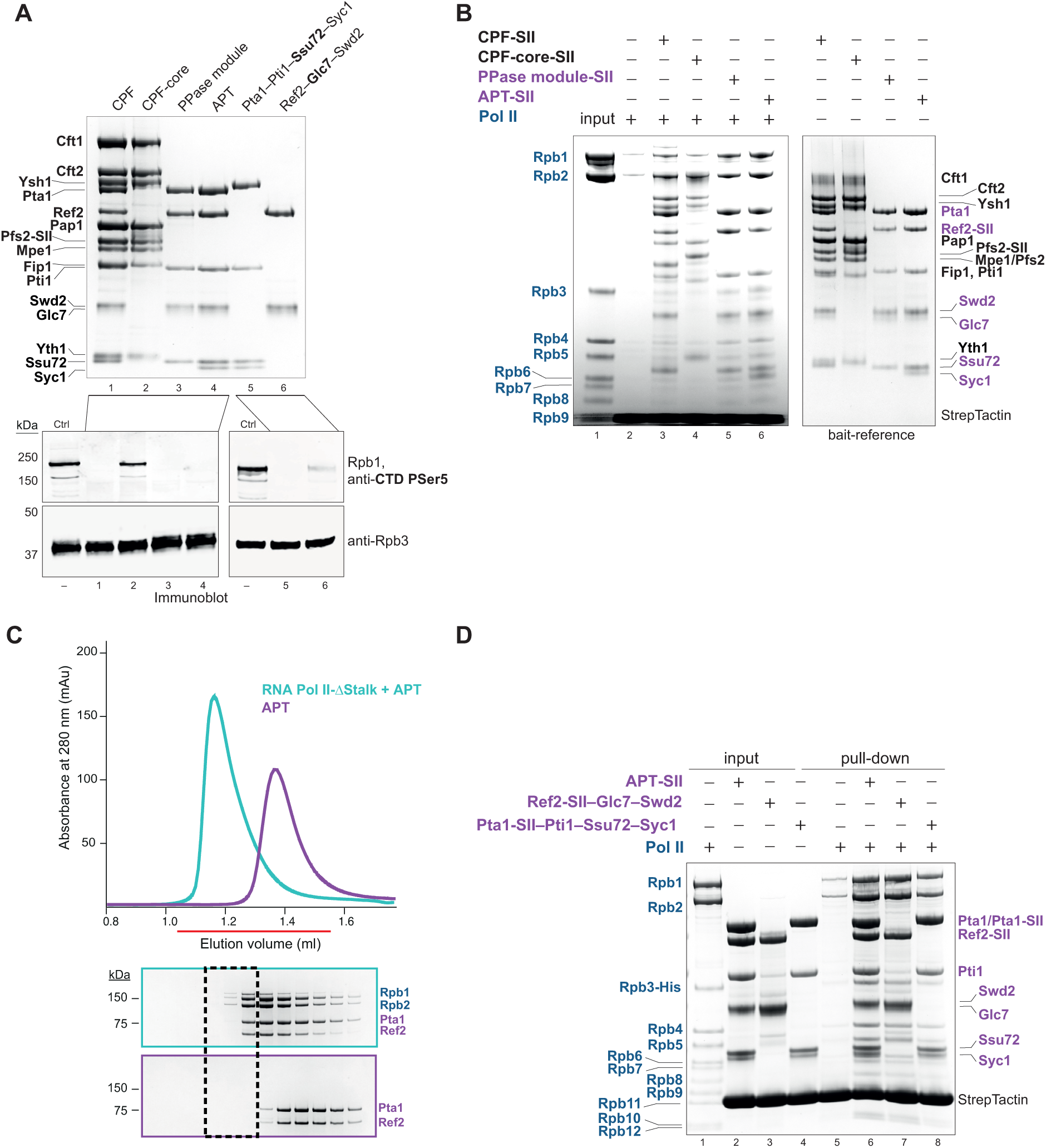
Characterization of the interaction between CPF and Pol II. (**A**) SDS-PAGE of CPF, APT and CPF subcomplexes. CPF-core includes the nuclease and phosphatase modules; PPase is phosphatase module. (Bottom) *In vitro* dephosphorylation assay of hyperphosphorylated Pol II with CPF complexes and APT, monitored by immunoblotting with an antibody against phosphorylated Ser5 (P-Ser5) CTD residues (3E8 antibody). Anti-Rpb3 (1Y26 antibody) was used as a loading control. Dephosphorylation assays against P-Tyr1 were not reproducible for recombinant proteins, possibly because this activity is affected by the CTD phosphorylation pattern, or due to variability in the 3D12 antibody. (**B**) Pulldown assay of untagged Pol II with SII-tagged CPF, CPF-core, phosphatase module (PPase) or APT immobilized on StrepTactin beads. SII indicates StrepII-tagged proteins. A gel with the bait proteins obtained from a different preparation is shown on the right as reference for comparison with lanes 2-6. Pol II subunits are labelled in blue (left); APT and PPase module subunits are labelled in purple and the remaining CPF subunits are in black (right). (**C**) Analytical size exclusion chromatography analysis of APT with Pol II- stalk. The red line denotes the fractions loaded on the SDS-PAGE below. The dashed black box highlights the migration position of the complex. The uncropped SDS-PAGE is in Supplementary Data 1D. (**D**) Pulldown assay of Pol II using StrepII (SII)-tagged APT (Ref2-SII), Ref2-SII–Glc7–Swd2, or Pta1-SII–Pti1–Ssu72–Syc1 immobilized on StrepTactin beads. Input and bound proteins were analyzed on SDS-PAGE. APT subunits are labelled in purple (right); Pol II subunits are labelled in blue (left). By following the Rpb3-His band, the Ssu72-complex interacts the most weakly with Pol II, compared to APT and the Glc7-subcomplex.

**Figure S2.**
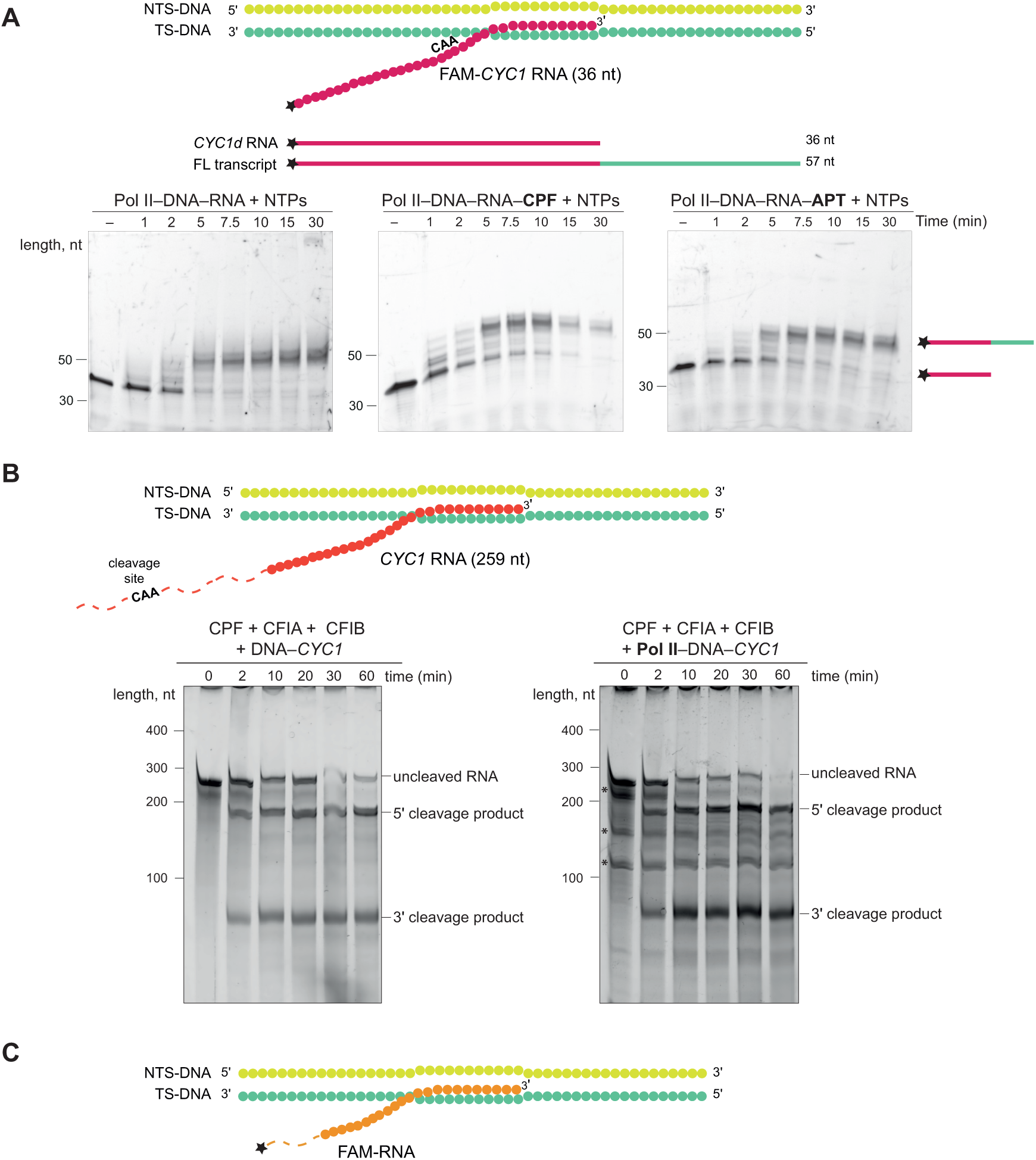
Functional interactions between CPF or APT and Pol II. (**A**) Promoter-independent transcription assay where Pol II was incubated with a DNA–RNA scaffold (NTS-DNA_A, TS-DNA_A and *CYC1d*-RNA, Table S2) and a mixture of nucleotide triphosphates (NTPs), with or without APT or CPF. NTS, non-template strand; TS, template strand; FL, full-length; *CYC1*, pre-mRNA substrate containing the CAA cleavage site (*CYC1d* fragment from Hill *et al*., 2019). Samples were collected at the indicated time points after the addition of NTPs. RNA extension was monitored on 20% denaturing urea-PAGE. The assays were performed in duplicate. (**B**) *In vitro* RNA cleavage assays of a 259-nt *CYC1* RNA annealed to the template strand DNA (TS-DNA_B, Table S2) (left) or annealed to an artificial transcription bubble (TS-DNA_B and NTS-DNA_D, Table S2) assembled with Pol II (right). The CAA cleavage site is depicted on the RNA and is 80 nucleotides away from the annealed sequence. Samples were collected at the indicated time points and cleavage products were analyzed on a 15% denaturing urea-PAGE. The asterisks on the right gel indicate degradation products of the RNA substrate prior to cleavage, likely resulting from the assembly procedure. The assays were repeated three times, and representative gels are shown. (**C**) Cartoon of the DNA–RNA hybrid mounted on Pol II for cryoEM analysis in complex with APT (TS-DNA_C, NTS-DNA_C and snR47 RNA, Table S2) or Ref2–Glc7–Swd2 (TS-DNA_D, NTS-DNA_D and RNA_D, Table S2). The latter RNA was based on (Ehara et al., 2017).

**Figure S3.**
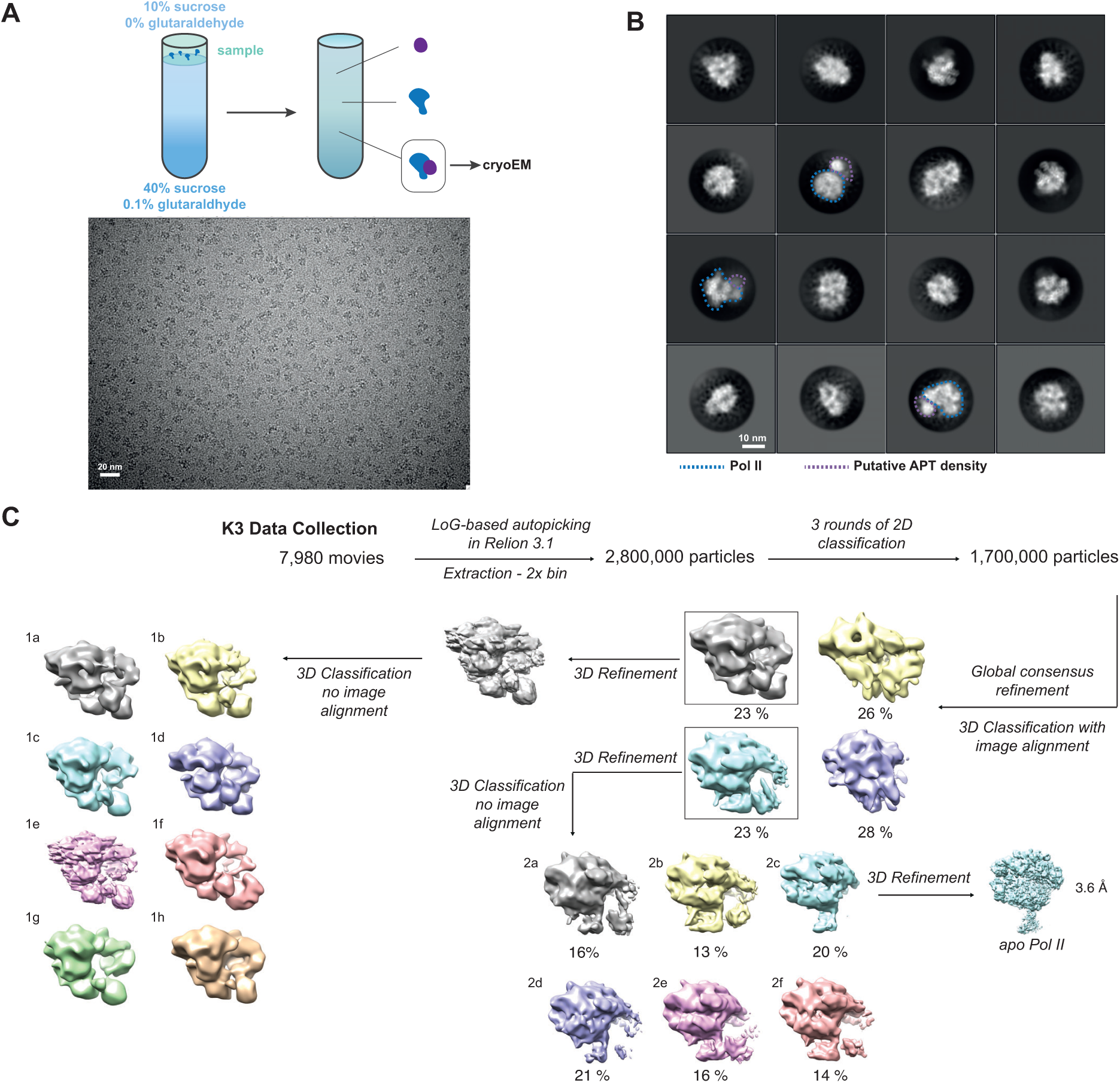
CryoEM analysis on DNA–RNA-loaded Pol II–APT prepared by GraFix. (**A**) (Top) Schematic of the GraFix protocol used for the crosslinking of Pol II–APT on a DNA–RNA scaffold. The gradient of sucrose and glutaraldehyde are indicated. The Pol II–APT mixture was applied on the top of the gradient (left). Blue and purple cartoons are Pol II and APT, respectively. On the right, the distribution across the gradient of APT, Pol II and their complex that was vitrified on grids (highlighted). (Bottom) Representative cryoEM micrograph collected on a Gatan K3 detector at a magnification corresponding to 1.06 Å/pixel. (**B**) 2D-class averages of the cross-linked Pol II–APT complex. Pol II core is highlighted in blue, and the putative APT density in purple. (**C**) Processing pipeline for the Pol II–APT data. Percentages represent the number of particles in each 3D-class, relative to the total particles from the previous step. Model 2a is shown in Figure 2A.

**Figure S4.**
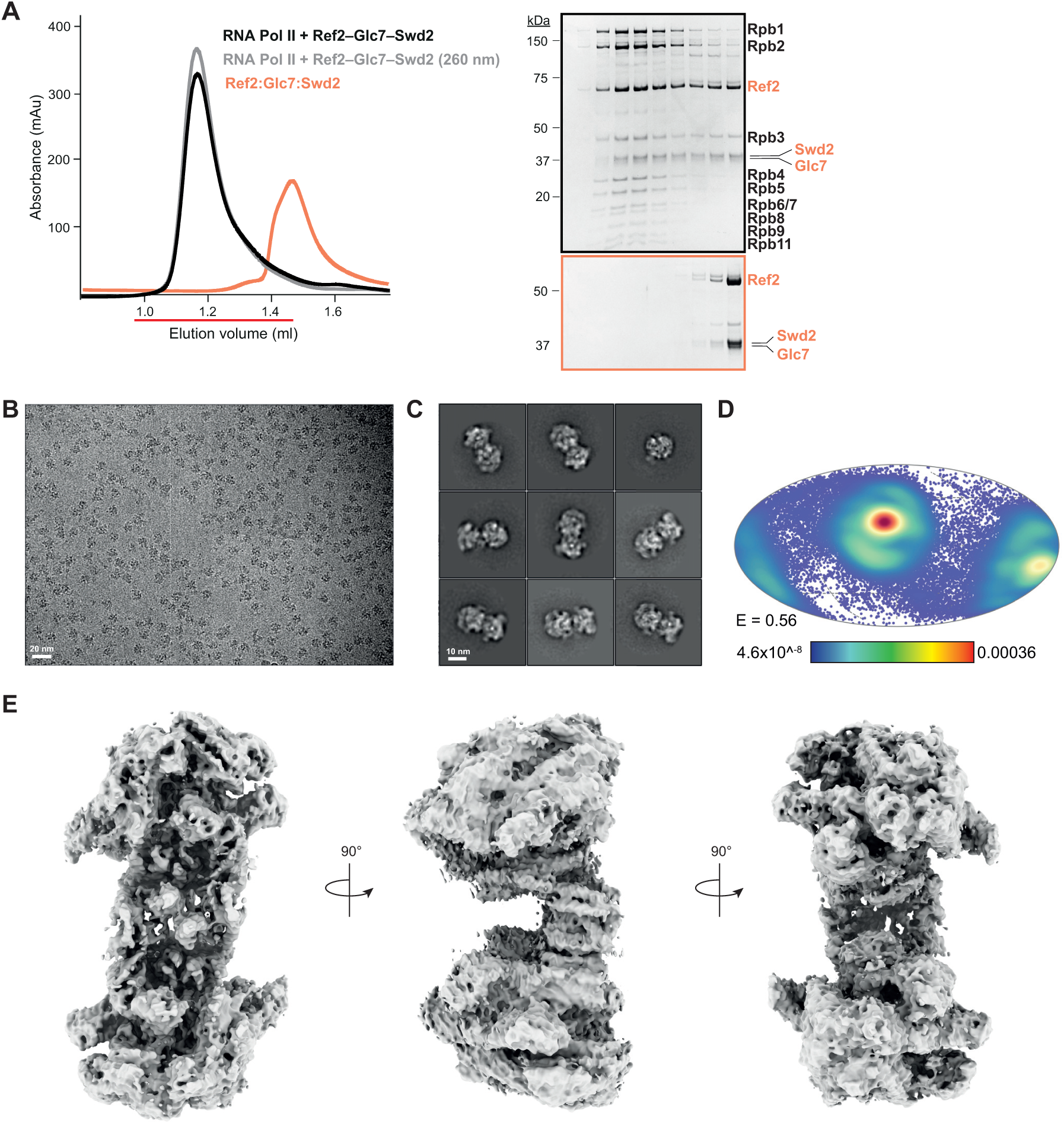
CryoEM analysis of Pol II bound to Ref2–Glc7–Swd2. (**A**) Size exclusion chromatography of Ref2–Glc7–Swd2 with Pol II in the presence of a DNA–RNA scaffold. The absorbance is at 280 nm except the grey trace which is at 260 nm. SDS-PAGE of selected fractions is shown on the right. The uncropped gels are shown in Supplementary Data 1C. (**B**) Representative cryoEM micrograph of the complex prepared under native conditions (not crosslinked). Data were collected on a Gatan K3 detector at a magnification corresponding to 0.83 Å/pixel. (**C**) Selected 2D class-averages from data collected at a 40° tilt angle. (**D**) Projection plot of orientation distribution of particles contributing to the final Pol II map shown in Figure 2D and Figure S4E. The red to blue color range denotes highest to lowest number of particles. E = cryoEF efficiency (Naydenova & Russo, 2017). (**E**) 3D reconstruction of the intact Pol II homodimer before focused refinement.

**Figure S5.**
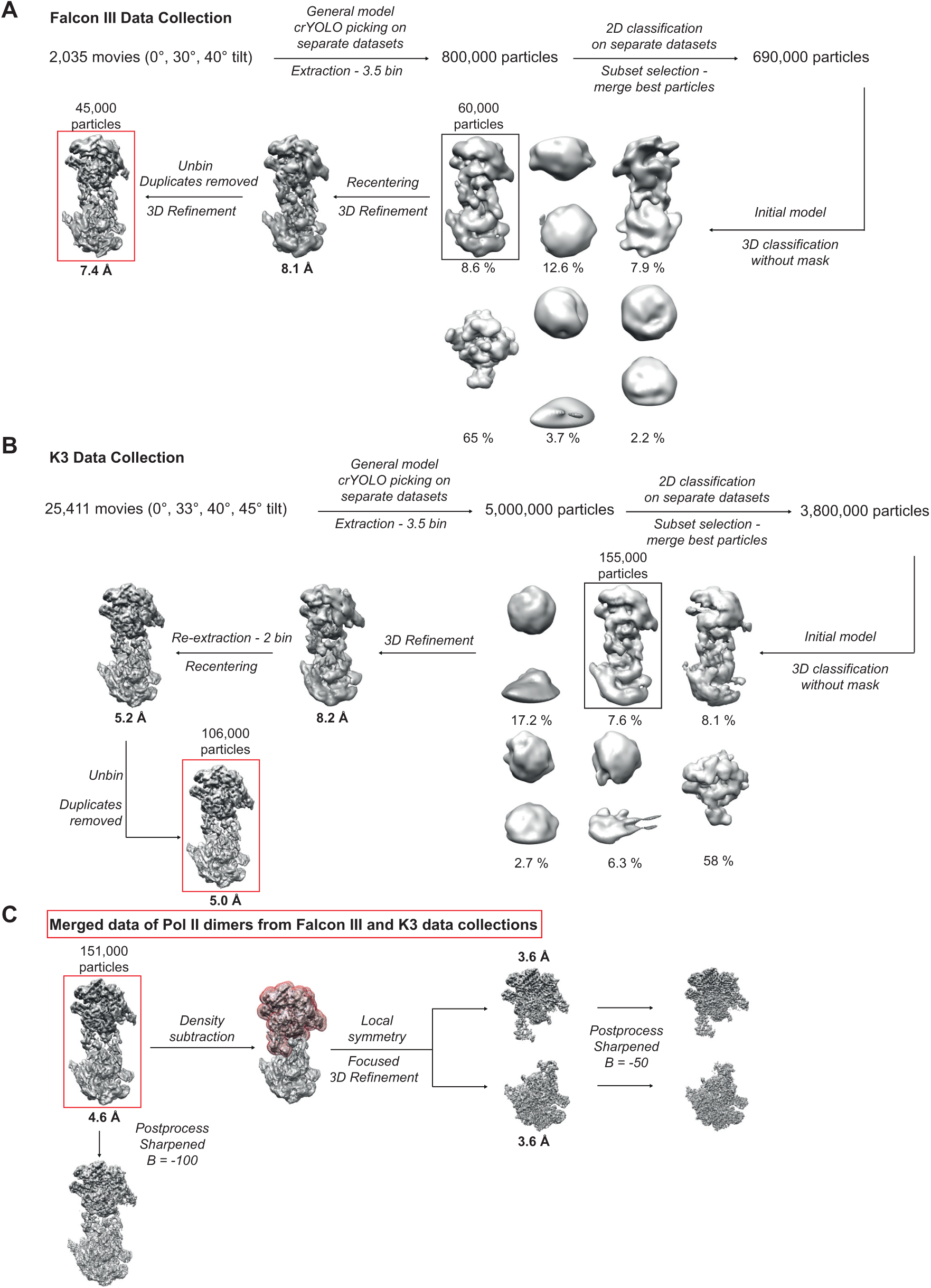
CryoEM processing pipeline of Pol II in complex with Ref2–Glc7–Swd2. (**A-B**) Flowchart of cryoEM processing for data the collected with or without stage tilt on a Falcon III (**A**) or a Gatan K3 (**B**) detector. Percentages represent the number of particles in each 3D-class, relative to the total particles from the previous step. (**C**) Particles from the final 3D reconstructions shown in panels **A** and **B** were merged and 3D refined (red box). The mask around the top monomer of the Pol II dimer map (red mesh) was used for signal subtraction. Maps were flipped on the Z-axis to match the correct handedness before rigid body fitting of two copies of monomeric Pol II (PDB 5C4X).

**Figure S6.**
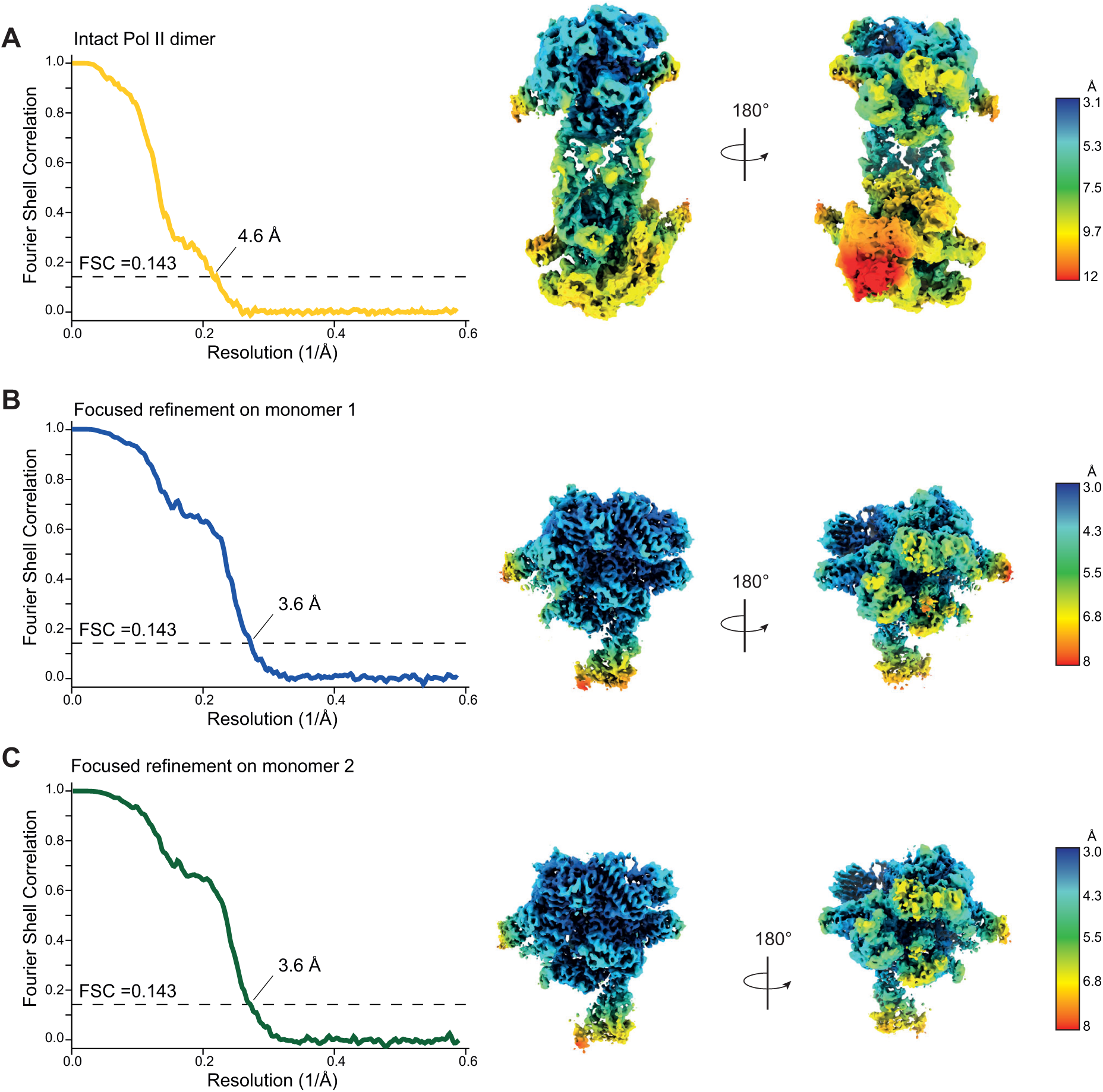
Pol II dimer cryoEM map validation and local resolution. (**A-C**) Fourier-shell correlation (FSC) curves of the map of the intact Pol II dimer (**A**), or the maps of monomer 1 (**B**) or monomer 2 (**C**) after focused refinement. Resolution was estimated according to the gold-standard FSC at 0.143. On the right, local resolutions estimated in Relion (v3.1) were plotted on the final sharpened maps.

**Figure S7.**
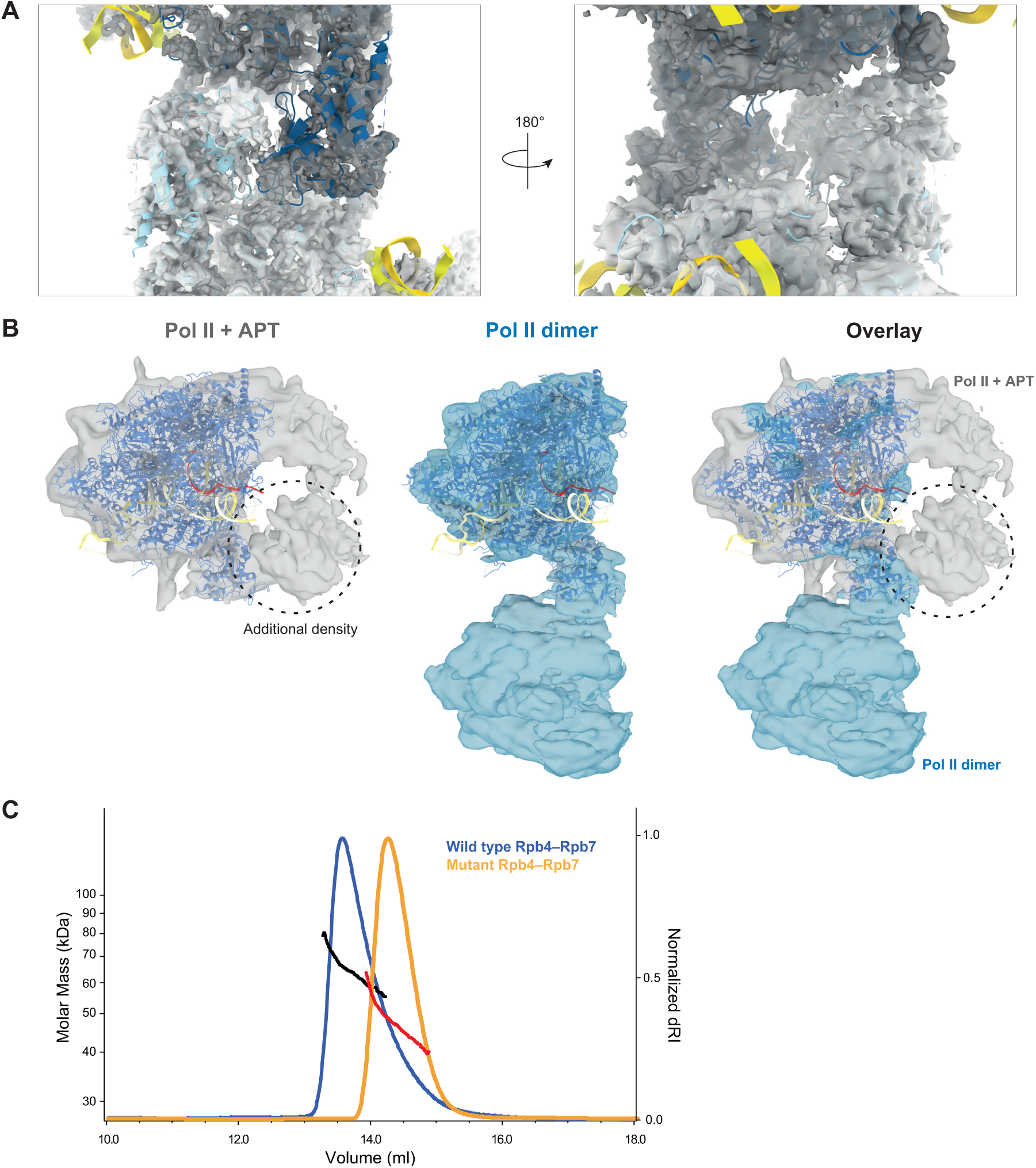
Validation and biophysics of the Pol II dimer interface. (**A**) Close-up view of the Pol II dimerization interface. The EM density is displayed as a transparent surface (grey). Fitted Pol II monomers 1 and 2 are shown in blue and cyan cartoon, respectively. The DNA–RNA hybrid is in yellow. The resolution of the stalk was not sufficient to confidently re-model it. (**B**) Comparison and overlay between the cryoEM map of crosslinked Pol II–APT (grey) and of the Pol II dimer (blue). The dashed circle indicates the putative APT density, which does not clash with the second monomer of the Pol II dimer. (**C**) SEC-MALS analysis of wild-type (blue) and mutant (yellow) Rpb4–Rpb7 (Pol II stalk) at 2 mg/ml. The blue and yellow traces represent the differential refractive index (dRI) of the samples, normalized to 1 and plotted against the y-axis on the right. The calculated molecular weights across the traces are shown in dotted black (wild-type) or red (mutant) lines and plotted against the left y-axis as 10^3^ g/mol (kDa). The lower average molecular mass of mutant Rpb4–Rpb7 is consistent with a shift towards a monomeric species in a concentration-dependent monomer-dimer equilibrium.

**Figure S8.**
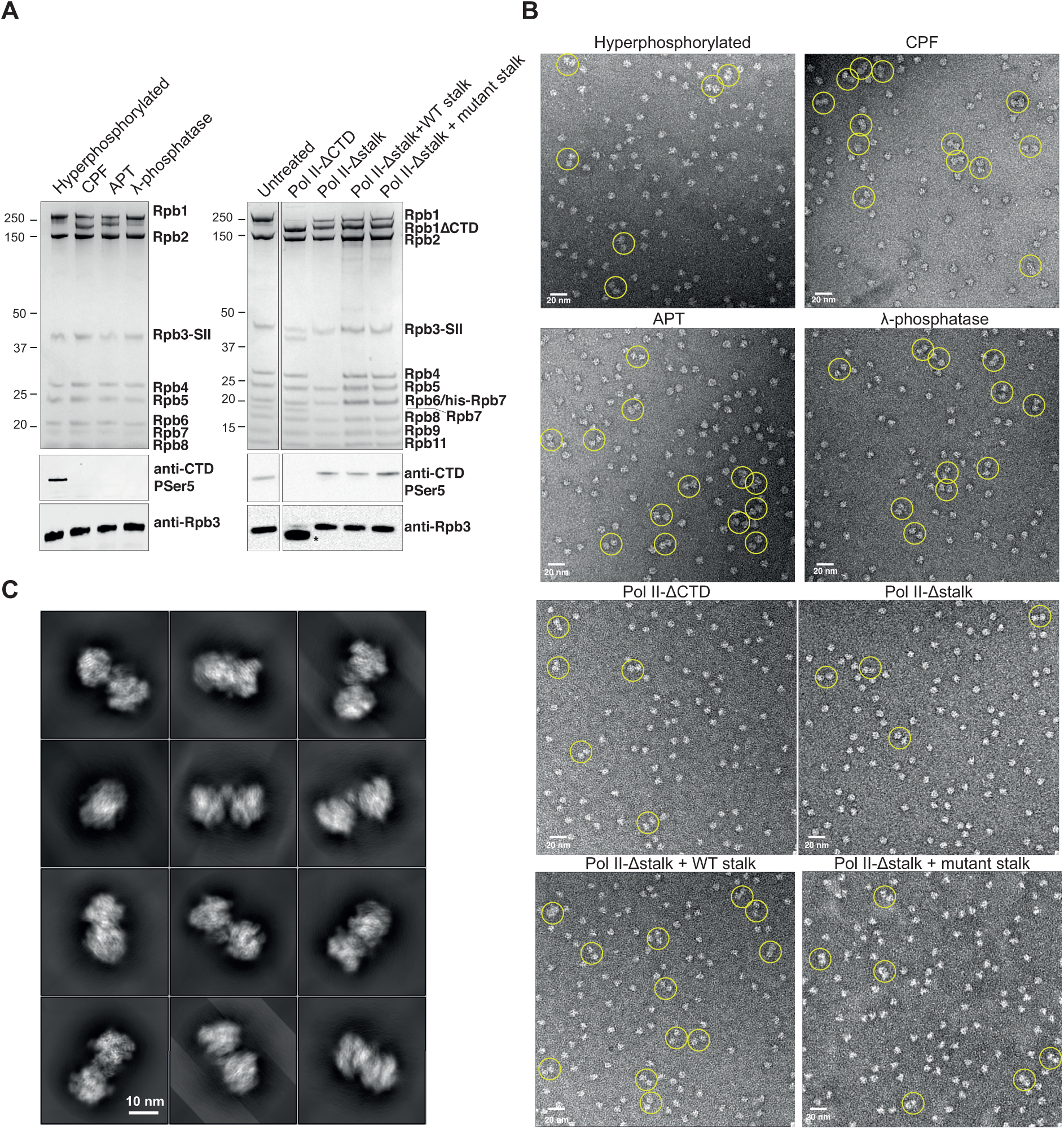
Negative Stain EM analysis on Pol II dimerization. (**A**) (Top) SDS-PAGE of the Pol II samples after different treatments used for the negative stain EM shown in panel B, analyzed in Figure 4C. (Bottom) Immunoblots against phosphorylated CTD Ser5 (antibody 3E8) to monitor the phosphorylation state for each condition. Anti-Rpb3 (antibody 1Y26) was used as a loading control. The shifted band marked with a black asterisk most likely resulted from non-specific recognition of a TEV cleavage site (within the Rpb3 tag) by 3C protease. (**B**) Representative negative stain EM micrographs of the various Pol II samples collected at a magnification of 21,000 X (2.53 Å/pixel). Data from 50-60 micrographs per condition were analyzed in Figure 4C. Particles counted as ‘dimeric’ Pol II are indicated with a yellow circle. (**C**) 2D projections of the Pol II homodimer cryoEM structure. The projections were used as a guide for manual inspection of ‘dimeric’ particles picked by Relion on the micrographs in **B**.

**Figure S9.**
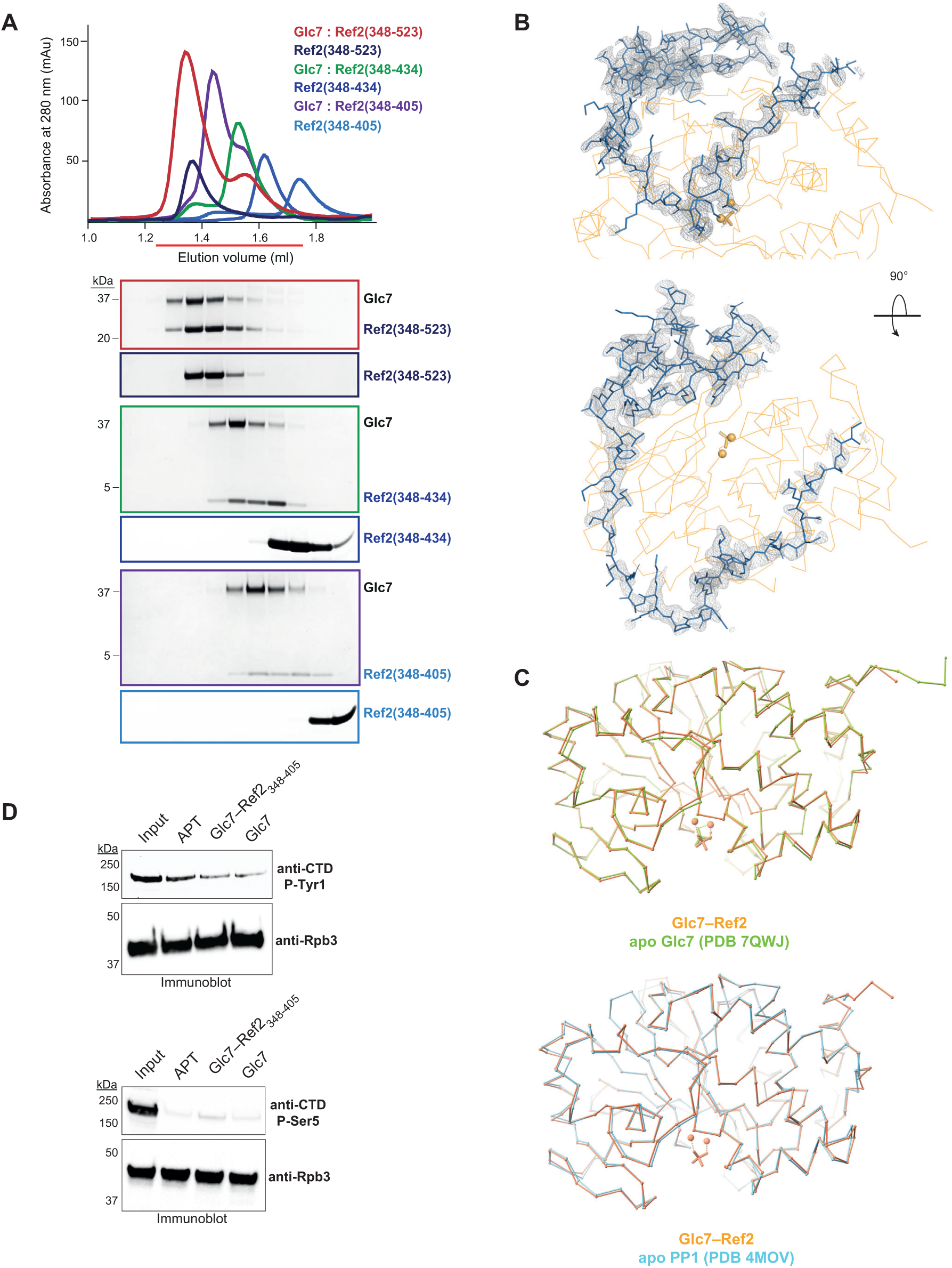
Structural analysis of Ref2 binding to Glc7. (**A**) Identification of the Glc7-binding region on Ref2 (fragments in shades of blue) by size exclusion chromatography. The elution profiles of the Ref2 truncations in isolation or in complex with Glc7 were overlayed, and the peak fractions were analyzed by SDS-PAGE. The outlines of the gels correspond to the colors of the chromatogram traces. (**B**) Two views of the Ref2_348-405_ peptide (blue sticks) built into the electron density obtained from X-ray crystallography. The grey mesh is an F_o_-F_c_ difference map displayed at 1.5 σ contour level. Glc7 is represented in ribbon (orange). (**C**) Overlay of the Ref2–Glc7 crystal structure (orange) with apo-Glc7 (green; PDB 7QWJ), and apo-PP1 (cyan; PDB 4MOV). The Ref2 peptide was omitted for clarity. (**D**) *In vitro* dephosphorylation assay of hyperphosphorylated Pol II comparing the activity of APT and isolated Glc7 to a Glc7–Ref2_348-405_ chimera lacking the CTD peptide. Immunoblots were performed with antibodies that recognize phosphorylated CTD Tyr1 (top) and phosphorylated CTD Ser5 (bottom) (antibodies 3D12 and 3E8, respectively). Anti-Rpb3 (1Y26) was used as loading control.

**Figure S10.**
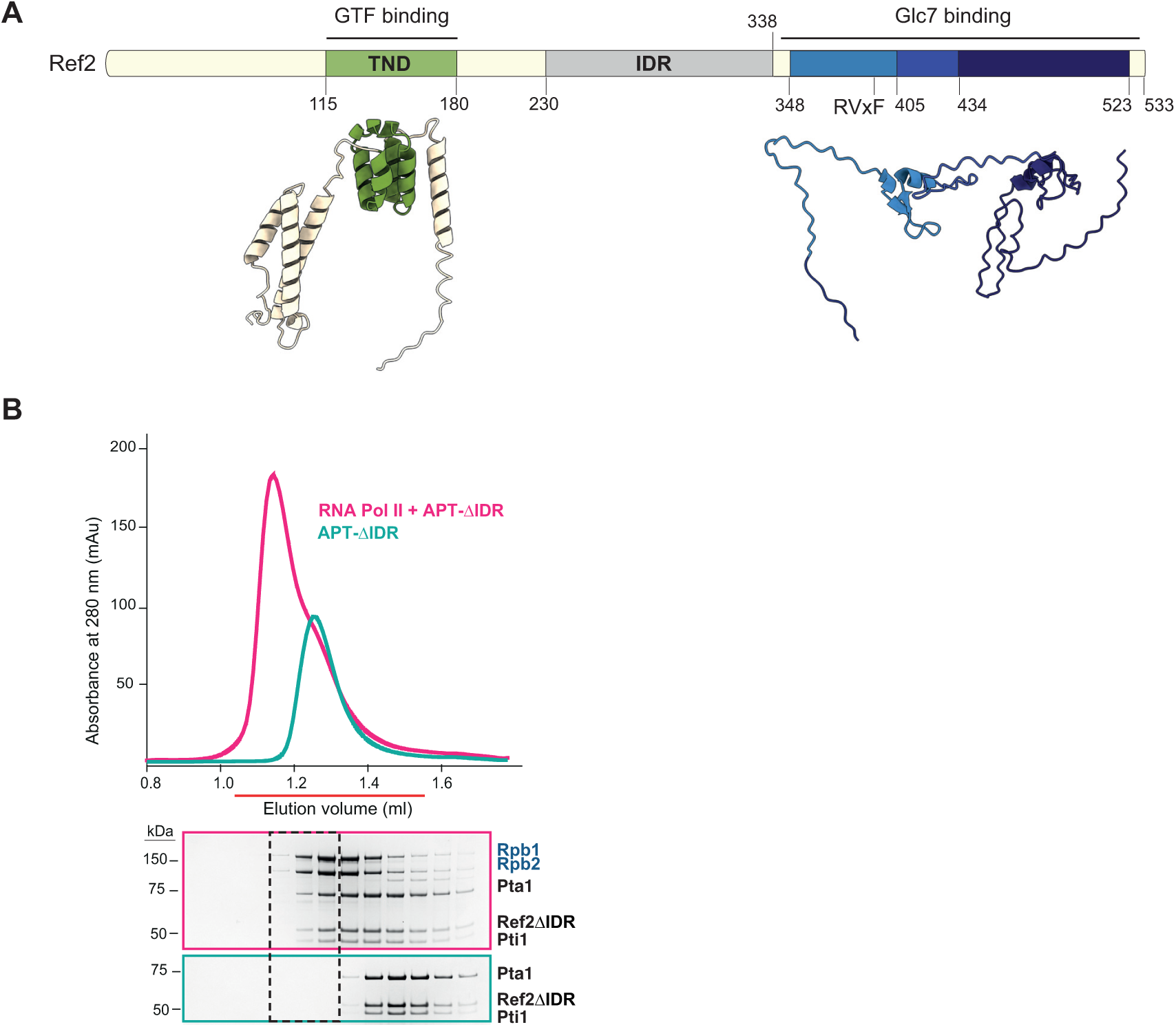
Characterization of the Ref2-mediated interaction with Pol II. (**A**) Domain diagram of yeast Ref2 (Uniprot P42073) with the AlphaFold2 structure prediction shown below in cartoon. The N-terminal region harbors a TFIIS N-terminal domain (TND) (green). A predicted intrinsically-disordered region (IDR) (grey) spans more than 100 residues in the middle of Ref2. The C-terminal region contains an RVxF motif that has been proposed to bind Glc7 *in vivo* (Nedea *et al*., 2008). The different shades of blue in the schematic and cartoon representation show the three fragments that were tested for Glc7 binding in Figure S9A. (**B**) Size exclusion chromatography of Pol II in complex with APT- IDR. The elution profile of the complex is in magenta, and the fractions indicated with a red line were loaded on SDS-PAGE. The black dashed line indicates the complex being formed. The outlines of the gels correspond to the colors of the chromatogram traces. The uncropped SDS-PAGE is in Supplementary Data 1E.

**Figure S11.**
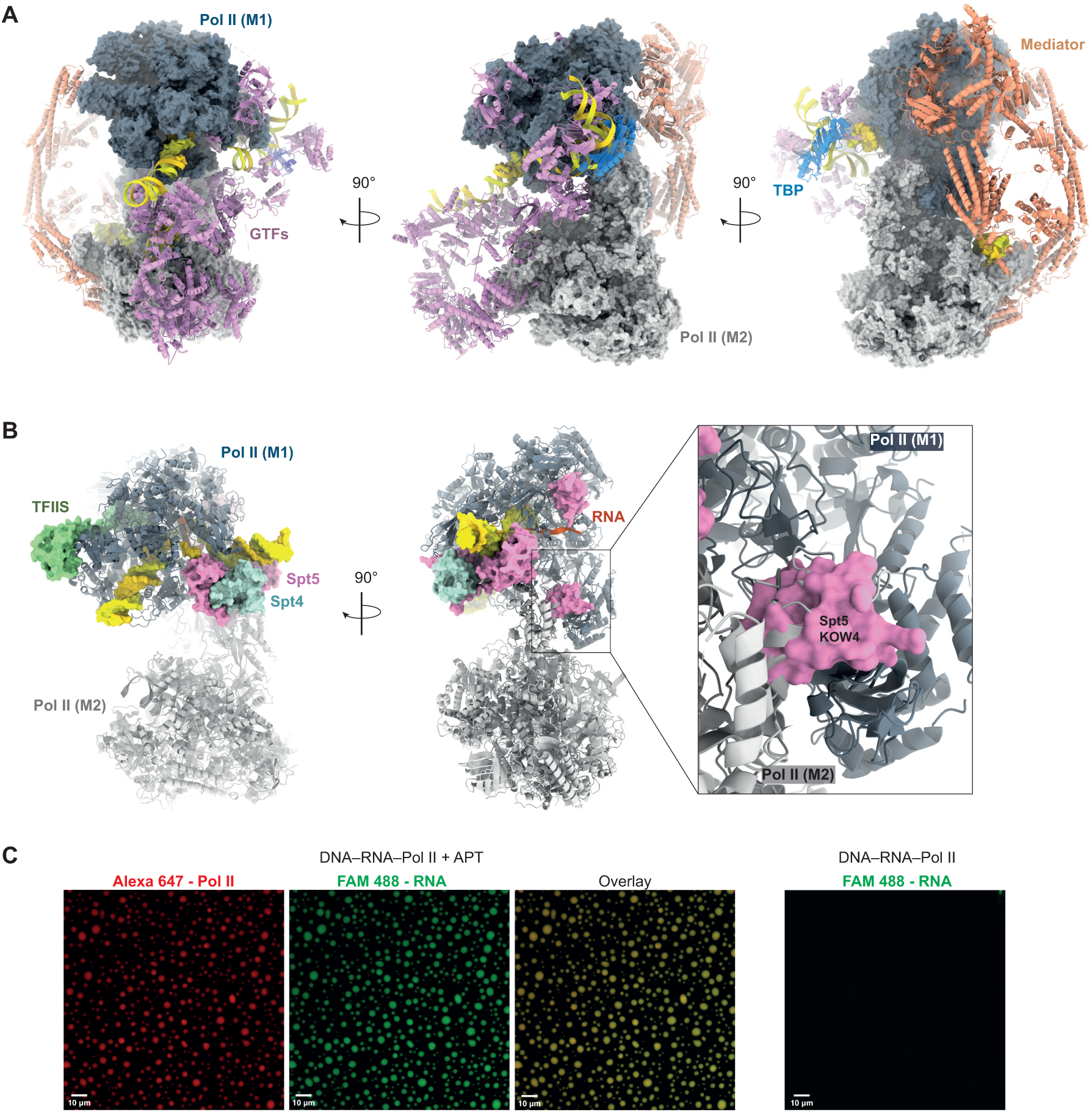
Overlay of the yeast Pol II dimer with other transcription complexes. (**A**) Views of the structural superposition between the yeast Pol II dimer (monomer 1 and monomer 2 in grey surface) and the pre-initiation complex (PIC) (PDB 5OQM) in cartoon. Mediator subunits are shown in salmon, TATA-binding protein (TBP) in blue, general transcription factors (GTFs) in pink, and DNA is in yellow. Monomer 2 has major clashes with mediator and GTF subunits including TFIIB, TFIIE and TFIIH. (**B**) Superposition of dimeric yeast Pol II (in cartoon) with the elongation complex (EC) shown in surface (PDB 5XON). EC Pol II has been omitted from the figure for clarity. TFIIS is in green, Spt5 in pink and Spt4 in cyan. The inset on the right shows a clash between monomer 2 from the dimer and the KOW4 domain of Spt5. (**C**) Confocal microscopy images of 0.6 μM Pol II - Alexa Fluor 647 loaded with a DNA–RNA(FAM) scaffold (Ehara *et* al., 2017 and Figure S2C) in the presence or in the absence of 4 μM APT. The overlay shows the co-localization of Pol II with the loaded DNA–RNA scaffold. The control on the right is Pol II mounted with the DNA–RNA hybrid without APT. Condensate formation can only be observed in the presence of APT. The experiment was repeated twice, and representative confocal acquisitions are shown.

**Supplementary Data 1.**
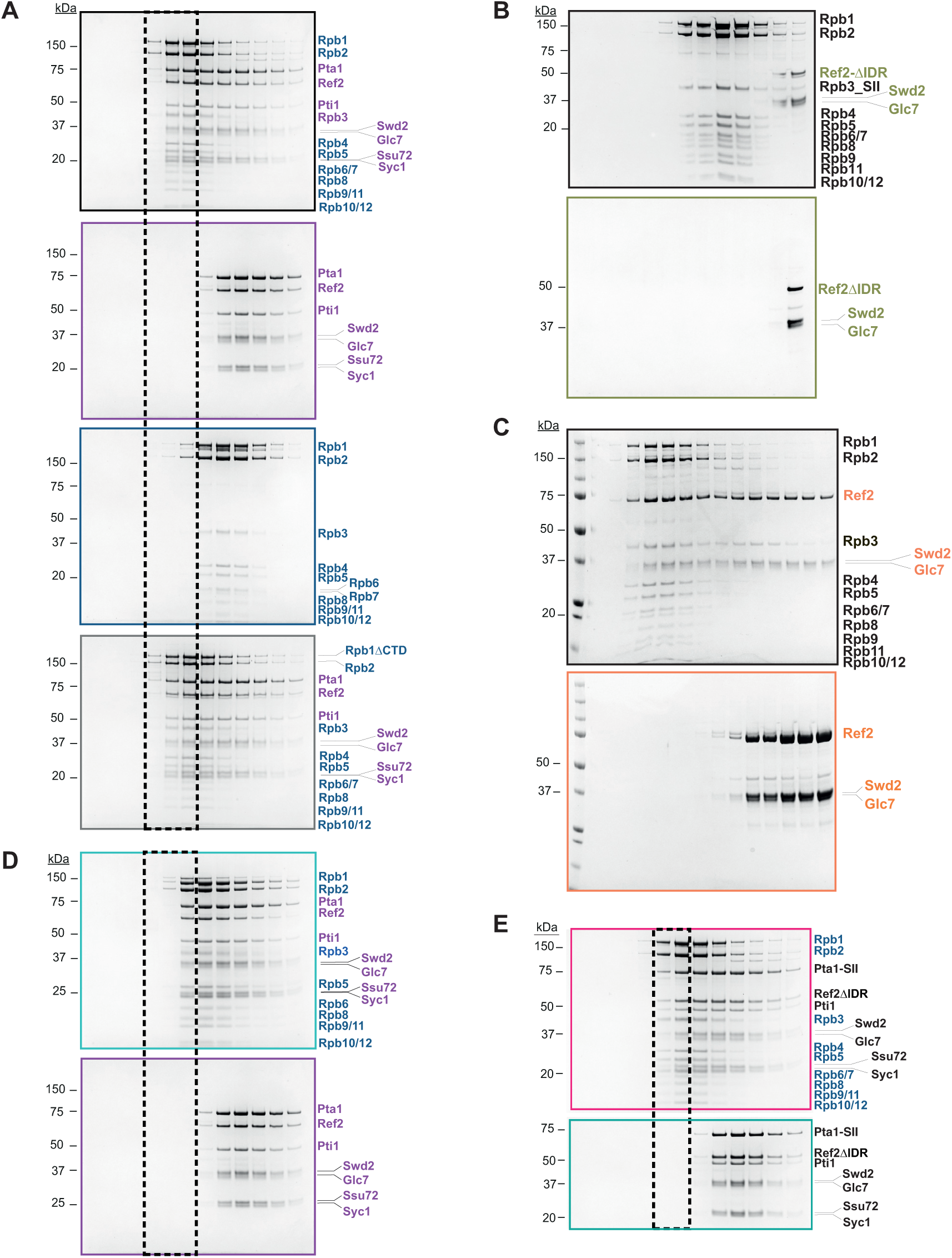
Uncropped SDS-PAGE panels across the study. **a**, Uncropped SDS-PAGEs of the eluted fractions from the SEC experiment in Figure 1B. Pol II subunits are labelled in blue, and APT in purple. **b**, Uncropped SDS-PAGEs of the eluted fractions from the SEC experiment in Figure 6B. Pol II subunits are in black, whereas Ref2- IDR–Glc7–Swd2 in green. **c**, Uncropped SDS-PAGEs of the eluted fractions from the SEC experiment in Figure S4A. Pol II subunits are in black, whereas Ref2–Glc7–Swd2 in orange. **d**, Uncropped SDS-PAGEs of the eluted fractions from the SEC experiment in Figure S1C. Pol II subunits are in blue, APT subunits in purple. **e**, Uncropped SDS-PAGEs of the eluted fractions from the SEC experiment in Figure S10B. Pol II subunits are in blue, APT subunits in black.

**Table S1.** List of yeast strains with genotypes.

| **Strain** | **Name** | **Genotype** | **Source** |
| --- | --- | --- | --- |
| *RPB3-His6-Bio*<br>*wt* | *BJ5464* | *MATa ura3-52 trp1 leu2-delta1 his3-delta200 pep4::HIS3 prb1-delta1.6R can1 GAL RPB3-His6-BIO::URA3* | Kireeva et al., (2000b)<br>J Biol Chem |
| *rpb4Δ* | *OCSC2065* | *MATa, ade2-1, his 3-11,15, leu2-3,112, trp1-1, ura3-1, rpb4::HIS3* | Allepuz-Fuster et al.,<br>2014 |
| *RPB3-TEV-His6-SII*<br>*rpb4Δ* | *MCY101* | *MATa, ade2-1, his 3-11,15, leu2-3,112, trp1-1, ura3-1, rpb4::HIS3 RPB3-TEV-His6-SII* | This study |
| *wt*<br>(*protease*<br>*deficient*) | *JWY104* | *MATalpha pra1-1 prb1-1 prc1-1 cps1-3 ura3delta5 leu2-3 his-* | Originally c13-ABYS-86 from Heinemeyer et al., (1991) EMBO J 10:555-562. |
| *RPB3-TEV-His6-SII*,<br>*Rpb1-3C-CTD*<br>*wt* | *MCY102* | *MATalpha pra1-1 prb1-1 prc1-1 cps1-3 ura3delta5 leu2-3 his-Rpb1-3C-CTD RPB3-TEV-His6-SII* | This study |

**Table S2.**
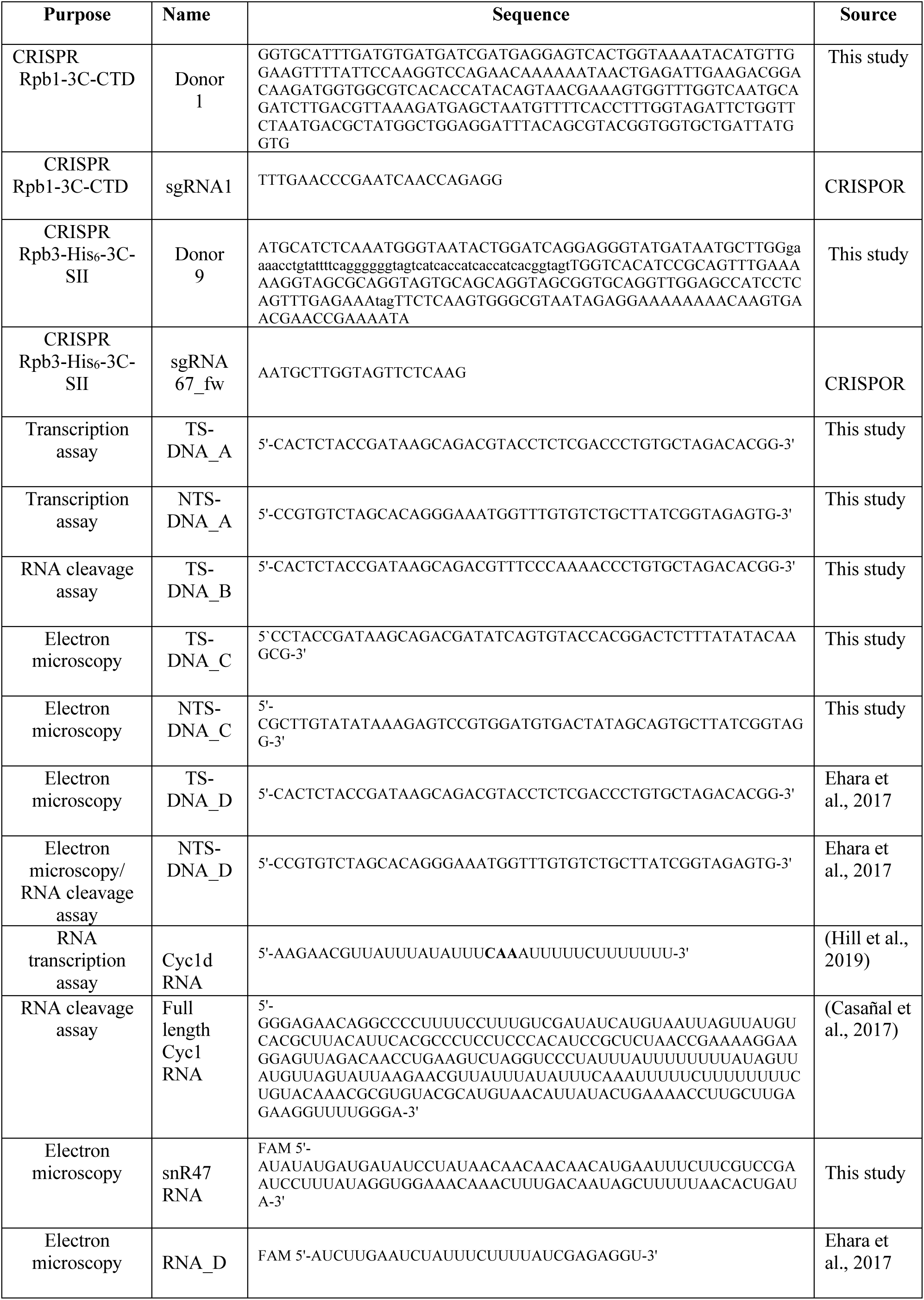
List of DNA and RNA oligonucleotides used across the study.

**Supplementary Movie 1**

Morph movie showing the relationship between the crystallographic Pol II dimer from Barnes *et al*., 2015, and the cryoEM Pol II dimer from this study. The two structures are related by a ∼15°rotation along the 2-fold symmetry axis.

